# Tracking and Measuring Local Protein Synthesis *In Vivo*

**DOI:** 10.1101/2021.06.27.450087

**Authors:** Ibrahim Kays, Brian E. Chen

## Abstract

Detecting when and how much a protein molecule is synthesized is important for understanding cell function, but current methods have poor cellular or temporal resolution or are destructive to cells (Dahm et al., 2008). Here, we developed a technique to detect and quantify subcellular protein synthesis events in real time *in vivo*. This Protein Translation Reporting (PTR) technique uses a genetic tag that produces a stoichiometric ratio of a small peptide portion of a split fluorescent protein and the protein of interest during protein synthesis. We show that the split fluorescent protein peptide can generate fluorescence within milliseconds upon binding the larger portion of the fluorescent protein, and that the fluorescence intensity is directly proportional to the number of molecules of the protein of interest synthesized. Using PTR, we tracked and measured protein synthesis events in single cells over time *in vivo*. We use split red fluorescent protein to detect multiple genes or alleles in single cells simultaneously. We also split a photoswitchable fluorescent protein to photoconvert the reconstituted fluorescent protein to a different channel and arbitrarily reset the time of detection of synthesis events, continually over time.

## Introduction

Understanding the relationship between genes and phenotypes is a central component of molecular and cell biology. Several analytic methods have been developed to profile gene expression at the level of mRNA or protein. The most common techniques for detecting protein synthesis are “pulse-chase” experiments to label previously translated proteins compared to newly synthesized proteins, or by measuring the production of a fluorescent protein or exogenous synthetic molecules as proxies for protein translation (Dahm et al., 2008; Hinz et al., 2013; Na et al., 2016; Wang et al., 2016; Yan et al., 2016). Genetically encoded fluorescent protein fusions have enabled the direct observation and tracking of intracellular proteins, however fusion proteins have been shown to affect protein properties and localization (Palmer and Freeman, 2004), affecting folding, timing, and function of both the protein of interest and the reporter (Zhao et al., 2008). In addition, constitutive fluorescence of common reporters may saturate measurement ranges over time, making the detection of small changes in signal intensity difficult, particularly *in vivo*. Genetically encoded luciferase approaches can provide high temporal or spatial resolution, but require repeated introduction of exogenous substrates necessary for the bioluminescence reaction, and this reaction can be inhibited by endogenously occurring molecules (Auld and Inglese, 2004; Ling et al., 2012; Na et al., 2016). The concentration of this substrate, when provided in the extracellular medium or by injection into animals, must be maintained throughout the experiment in order to produce accurate and consistent results, and several instability issues have been reported since early uses of luciferases (Craig et al., 1991; Morse and Tannous, 2012; Xu et al., 2016).

Approaches such as SunTag (Tanenbaum et al., 2014), SINAPS (Wu et al., 2016), and HA-based tagging (Zhao et al., 2019) employ antibody-based fragments to recruit and concentrate multiple fluorophore molecules. These approaches have enabled the visualization of the dynamics of nascent peptide chain production and tracking single molecules for extended periods of time *in vivo*. However, the recruitment and eventual fusion of several fluorophore molecules has been shown to alter several properties of the proteins of interest, such as size, charge, solubility, diffusion, turnover and localization (Snapp, 2005; Yang et al., 2016).

Furthermore, tagging endogenous loci with repeat sequences that are synthetic can result in dysregulation in the expression and production of the protein of interest (Yang et al., 2016). Finally, while these approaches enable the visualization of low-abundance proteins by multiplying the fluorescent signal, quantifying the production of secreted or short-lived proteins such as antibodies, transcription factors, or circadian clock proteins may not be possible with these techniques (Cohen et al., 2014; Corish and Tyler-Smith, 1999; Landgraf et al., 2012; Lepore et al., 2019).

We have recently developed a method called Protein Quantitation Ratioing (PQR) to quantify endogenous protein translation in single cells *in vivo* using a fluorescent reporter (Lo et al., 2015). The PQR sequence allows for an equimolar separation of an upstream protein of interest and a downstream reporter protein, all contained within a single strand of RNA. When a fluorescent protein reporter is separated from the protein of interest, the number of fluorescent molecules produced are proportional to the number of protein molecules of interest produced, and thus the fluorescence intensity is proportional to the target protein concentration. This PQR technology is essentially a quantitative protein translation reporter because the stoichiometric production of the fluorescent reporter molecule occurs at the protein translation step. However, the slow time scales (minutes to hours) for fluorescent proteins to fold and mature severely limit their use in reporting protein synthesis (Iizuka et al., 2011; Shaner et al., 2005), due to the speed of molecule diffusion at small, subcellular length scales. A typical protein might diffuse across a 20 μm cell in 40 seconds, but will diffuse 350 μm in the ten minutes it may require to fold and mature. The output of any protein synthesis reporter must occur within milliseconds to accurately detect synthesis events within the micrometer length scale or risk degrading spatial and temporal resolution.

## Results

In an effort to overcome the folding and maturation issues of common fluorescent proteins, we have generated Protein Translation Reporters (PTRs), which use a small peptide that is co-translated with the protein of interest, and will bind and activate a fluorescent reporter after its co-translation (**Figure 1a**). The peptide is based on the complementation of a split green fluorescent protein (GFP) into two non-fluorescent parts (Feinberg et al., 2008; Kerppola, 2006): a larger portion containing ten of the eleven strands of the GFP beta-barrel structure (GFP1-10), and the eleventh strand of the beta barrel comprised of only 16 residues (GFP11). High levels of expression of GFP1-10 in a cell allow for folding and chromophore maturation to occur independently and prior to protein translation reporting. Using split GFP components as Protein Translation Reporters (PTRs) to monitor protein translation events offers a number of advantages over probe or antibody-based approaches. For example, PTRs are genetically encoded and signals are fluorescence-based, which minimizes the cell or animal invasiveness associated with detecting protein translation. The untagged protein of interest remains free in its native form after synthesis, which ensures proper localization, secretion or post-translational modification. In addition, the fast reconstitution of the reporter immediately after translation would provide exceptional spatial and temporal resolution that can be used to localize protein translation events in subcellular compartments such as the endoplasmic reticulum or neuronal dendrites. Such an approach would open the door to direct, non-invasive and long-term observation of endogenous local protein synthesis in neurons, which is key to understanding the distal processes that both maintain cellular homeostasis and mediate plasticity.

**Figure 1.**
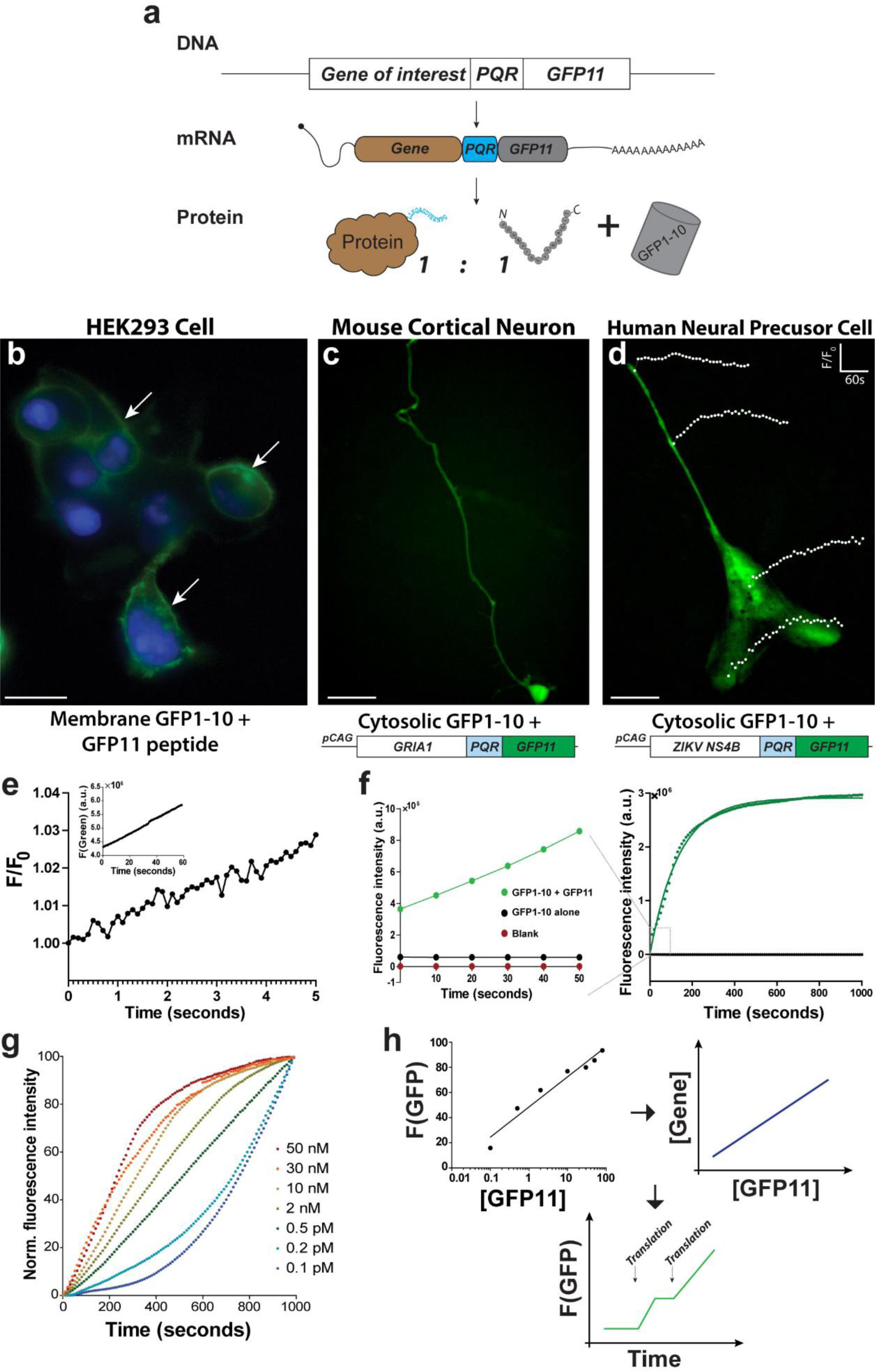
Stoichiometric production of GFP11 reporter using PQR allows instantaneous detection of protein translation. **a**, Insertion of a *Protein Quantitation Reporter* (PQR) between a split GFP11 reporter (GFP11) and a gene of interest creates a polycistronic mRNA for co-transcription and co-translation of GFP11 and the gene of interest. The *PQR* construct allows for one molecule of GFP11 to be synthesized for every one molecule synthesized of a protein of interest. In the presence of GFP1-10, the split GFP parts reconstitute and produce green fluorescence. **b**, Split GFP reporters can be expressed and reconstitute properly using PQRs. Membrane-tagged GFP1-10 was expressed on the extracellular surface of HEK293 cells and produced a bright green membranous fluorescent signal upon addition of 50 μM GFP11 peptide into the culture medium. **c, d**, Split GFP components expressed within the cytosol reconstituted to produce green fluorescence signal in mouse cortical neurons (**c**) and human neural precursor cells (**d**) *in vitro*. Mouse cortical neurons co-expressed cytosolic GFP1-10 and the *GRIA1* gene (Glutamate Ionotropic Receptor AMPA Type Subunit 1) tagged with *PQR-GFP11* (**c, Supplementary Movie 2**). Human neural progenitor cells co-expressed the Zika virus NS4B protein with *PQR-GFP11* and GFP1-10 (**d, Supplementary Movie 3**). White traces in **d** are the changes in green fluorescence across different subcellular regions of interest, representing Zika virus NS4B local translation over time. **e, f**, Reconstitution of split GFP occurs within milliseconds *in vitro*. Fluorescence reconstitution kinetic traces of split GFP shows the linear phase of reconstitution over the first 5 seconds (**e**), and the first minute (**inset**). GFP1-10 and GFP11 were mixed at a 200:100 nmolar ratio at 37°C and the fluorescence intensity was recorded over time. Within the first second of reconstitution, fluorescence emission begins, and the fluorescence intensity of the reaction rose logarithmically and began to saturate after 5 minutes (**f**). The fluorescence increase against time was fit to a one-site model and the observed rate constant k_obs_ for 50 nM GFP11 was determined to be 0.00335 s^-1^, (*R*^2^ = 0.99, *p* < 0.001). **g**, GFP11 has high affinity for GFP1-10. Fluorescence reconstitution kinetic traces for varying amounts of GFP11 peptide mixed with an excess of GFP1-10 were fit with a one site binding model. The dissociation constant K_d_ of GFP11 is 481 ± 116 pM (*R*^2^ = 0.96, *p* < 0.05) for binding GFP1-10 and unlikely to dissociate. The association rate constant k_on_ is 7.6 × 10^5^ M^−1^ s^−1^. **h**, A series of linear relationships allows for PTR to quantify protein translation events. First, the fluorescence intensity of the reconstituted GFP is linearly dependent on the input concentration of GFP11 over two orders of magnitude (**left panel**, data taken from concentration curves). Second, the level of GFP11 production is proportional to the level of protein of interest production (**right panel**, schematic) due to PQR (Lo et al., 2015). Thus, the fluorescence intensity of GFP reconstitution (i.e., brightness) can be used to determine the moment and amount of synthesis of the protein of interest over time (**bottom panel**, schematic). Scale bars are 30 μm in **b** and **d,** and 60 μm in **c**.

To demonstrate our approach, we screened through several variants of GFP and found that some versions of split GFP (Do and Boxer, 2011; Feinberg et al., 2008; Kent and Boxer, 2011; Kim et al., 2011; Pedelacq et al., 2006; Yamagata and Sanes, 2012) would not always express properly in cells and could aggregate in inclusion bodies or misfold, so we screened through different modifications that added specific amino acids to the carboxy and amino terminus of GFP11 and GFP1-10, respectively (**Methods, Table 1**). Expression of GFP1-10 on the cell surface of HEK293 cells produced green fluorescence when exposed to extracellular GFP11 peptide (**Figure 1b**). Next, we genetically tagged the *GRIA1* gene (Glutamate Ionotropic Receptor AMPA Type Subunit 1) using *PQR-GFP11* (i.e., *GRIA1-PQR-GFP11*), and co-expressed this with cytosolic GFP1-10 in mouse cortical neurons using an Actin promoter. Co-expression of these two constructs produced bright green cytoplasmic fluorescence signals that were not sequestered into inclusion bodies or lysosomes (**Figure 1c**). We were able to image protein translation dynamics of the GLUA1 subunit of the AMPA receptor as changes in green fluorescence over time. We also tagged the Zika virus (*ZIKAV*) protein NS4B using *PQR-GFP11* and expressed this with GFP1-10 within human neural progenitor cells (hNPCs) (**Figure 1d**). Similar to the GLUA1 subunit, these protein synthesis dynamics varied across different sampling locations within the cell body and along neurites (**Figure 1d**), indicating local differences in protein production levels.

**Table 1:**
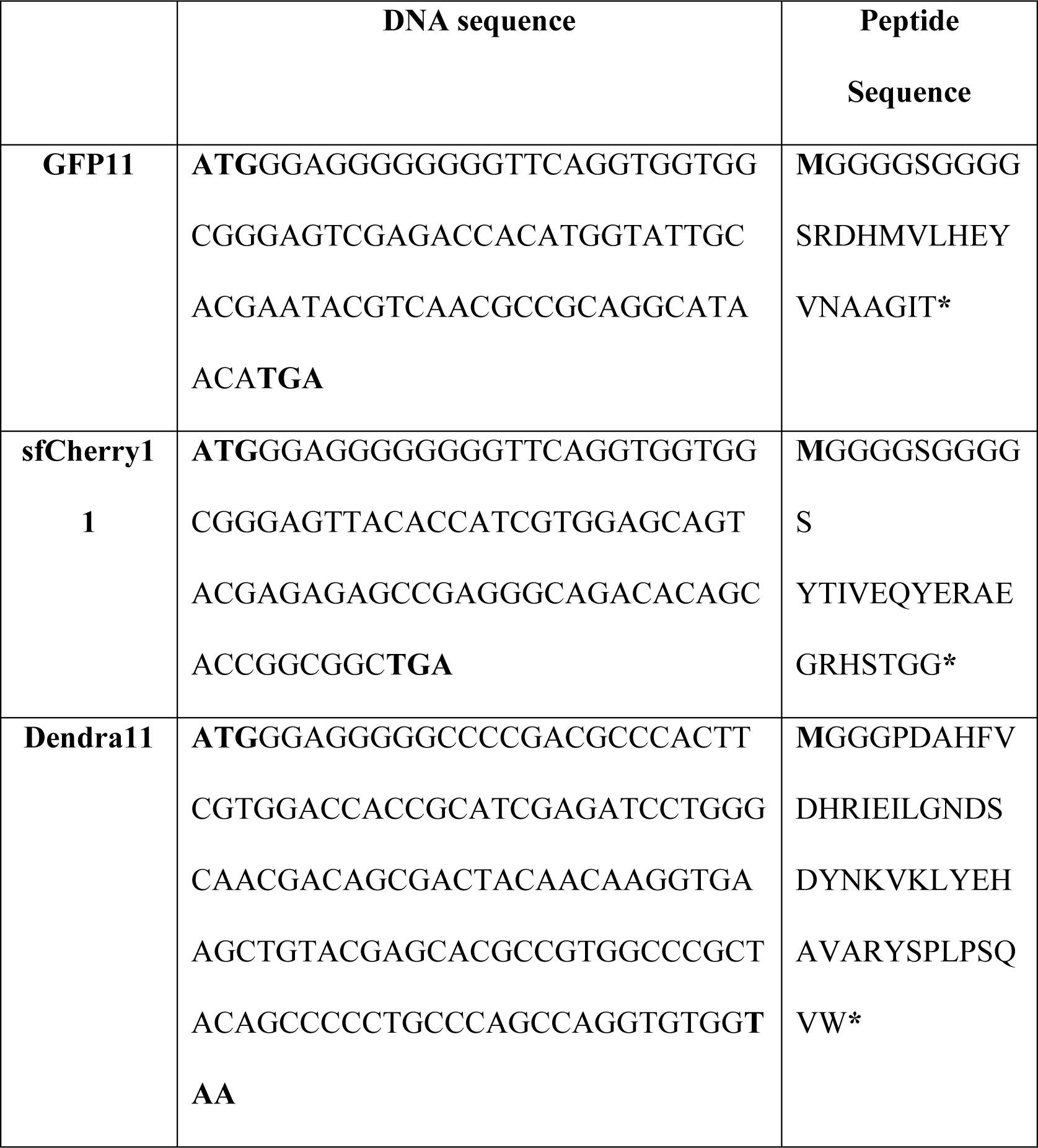
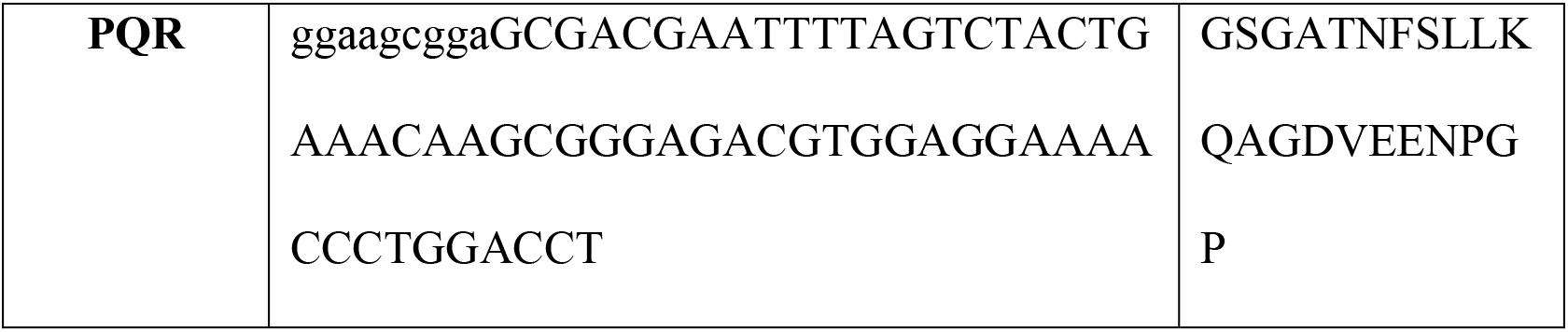
Sequences of PQR and PTR variants used.

To determine the speed of reconstitution of our modified split GFP, we combined purified GFP1-10 at 3 mM with varying concentrations of GFP11 and found that fluorescence was detected immediately upon mixing the two solutions (**Figure 1 e–g**). To obtain kinetic measurements of the reconstitution reaction at sub-second timescales, we used an ultrasensitive fluorescence spectrophotometer (**Methods**). A stopped-flow dispenser was used to mix and deliver known volumes of GFP1-10 and GFP11 proteins. This allowed us to establish a t_0_ for the moment split GFP components first interact. Green fluorescence was detected immediately upon mixing the two solutions (within 100ms) (**Figure 1e**), and the fluorescent intensity of the reconstitution reaction steadily increased throughout the recording window (∼60 seconds) (**Figure 1e, inset**). Fluorescence increase over time from the bimolecular reaction was modeled as pseudo-first order and fit to a one phase association curve to determine the observed rate constant (k_obs_) of GFP11 peptide binding for each concentration of GFP11 tested (**Figure 1f, g, Supplementary Figure 1, Methods**). For 50 nM GFP11 and excess GFP1-10, our k_obs_ is 0.003 s^-1^. Using these k_obs_, the association rate constant (k_on_) was calculated as 7.6 × 10^5^ M^−1^ s^−1^, which falls within the diffusion-controlled regime (Schreiber et al., 2009). To determine the dissociation constant (K_d_) of the GFP11 peptide, the fluorescence increase over time with varying GFP11 peptide concentrations was also fit to a one site binding model, and the K_d_ was 481 ± 116 pM (*R^2^* = 0.96, *p* < 0.05), demonstrating that GFP11 peptide binds GFP1-10 with very high affinity (Do and Boxer, 2011; Huang and Bystroff, 2009; Kent and Boxer, 2011). In the volume of a medium-size cell, the slower, lowest concentration reactions would generate tens of reconstituted molecules per second, and for the average protein synthesis hundreds per second (**Supplementary Figure 1**).

The fluorescence intensity resulting from the reconstitution of GFP1-10 and GFP11 peptides *in vitro* was linearly dependent on the concentration of the reactants over several orders of magnitude (**Figure 1h, top left**). In addition, expressing the GFP11 peptide using PQR linkers (**Figure 1a, c, d**) produces the stoichiometric relationship with the protein of interest (**Figure 1h, top right**) (Lo et al., 2015). Thus, the fluorescence intensity resulting from the translation of GFP11 is proportional to and can be used to quantify the level of translation of a protein of interest at the millisecond timescale (**Figure 1h, bottom**).

Does the genetic tag of *PQR* with *GFP11* produce a linear relationship with the protein of interest? We used the *Drosophila* Shaker potassium channel with a red fluorescent protein (RFP) embedded within the inactivation domain (Lo et al., 2015) to verify that the reconstituted GFP could quantify protein synthesis (**Figure 2a**). We recorded Shaker K+ currents from HEK293 cells expressing the *ShakerRFP* gene tagged with *PQR-GFP11* (*ShakerRFP-PQR-GFP11*) along with *GFP1-10* and confirmed that the PTR did not disrupt protein function (**Figure 2b**). RFP fluorescence intensity was correlated with reconstituted GFP with an *R^2^* = 0.72, *n* = 35, *p* < 0.05 (**Figure 2c**). We observed GFP puncta throughout the cell, indicating sites of local translation, possibly at ribosomes near the rough endoplasmic reticulum, as the Shaker potassium channel is being processed while the GFP11 diffuses away in the cytosol.

**Figure 2.**
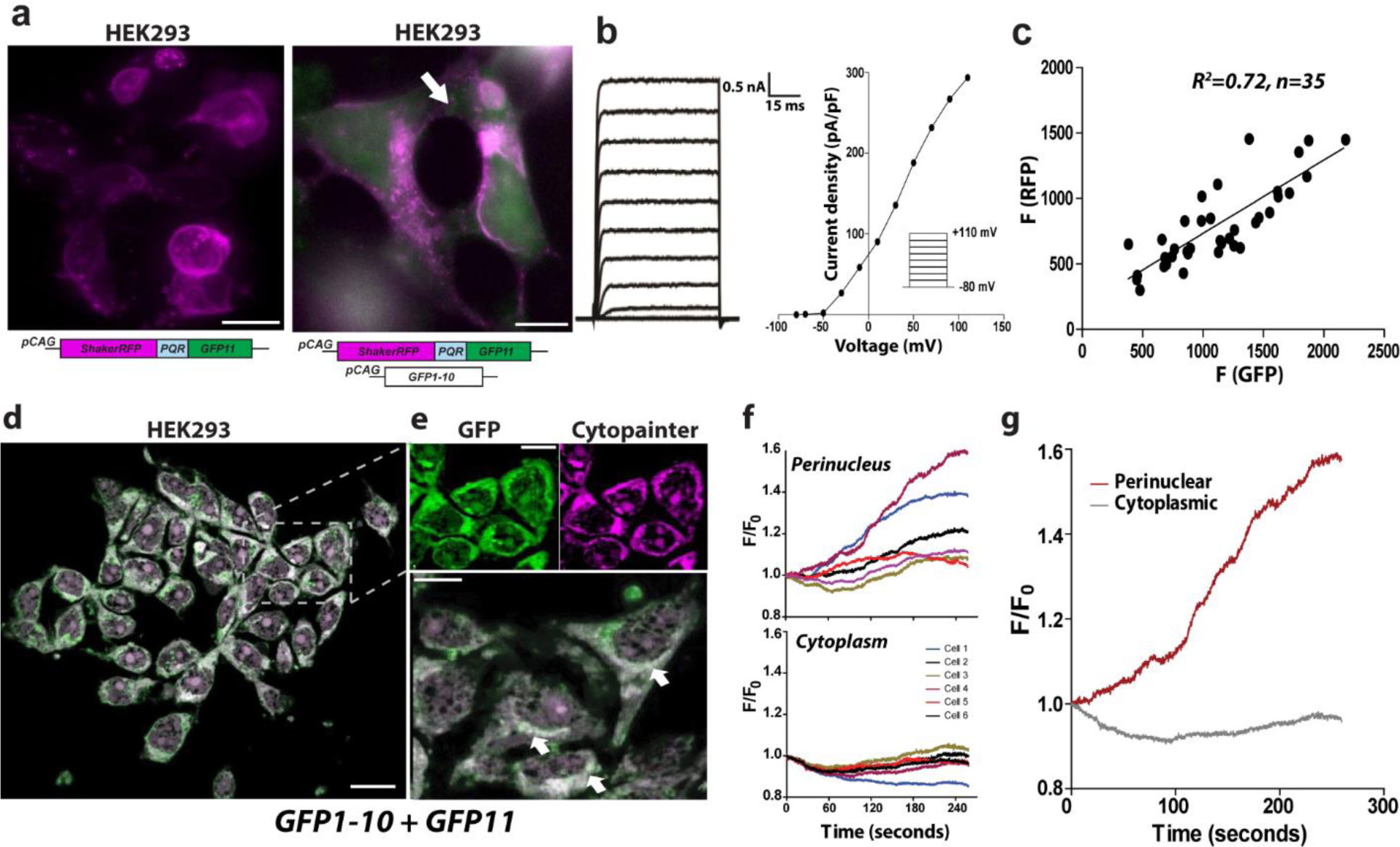
Direct observation of protein synthesis at ribosomes. **a-c**, Using PQR to co-translate GFP11 preserves the protein of interest’s localization and function. Expression of *ShakerRFP-PQR-GFP11* in HEK293 cells (**a**) produced red fluorescence at the cell membrane, indicating that the ShakerRFP potassium channel was processed and inserted correctly. With the co-expression of GFP1-10, the GFP11 reconstituted with GFP1-10 and produced cytoplasmic green fluorescence (arrow) that remained in the cytoplasm. Potassium conductances (**b**) in cells transfected with *ShakerRFP-PQR-GFP11* were consistent with previous Shaker currents (Lo et al., 2015). Green and red fluorescence intensities with linearly correlated (**c**), indicating that the level of GFP11 production is proportional to the level of ShakerRFP production. **d-e**, Reconstitution of split GFP11 occurs immediately after translation. HEK293 cells expressing split GFP constructs (**d**) were labeled with a red marker for endoplasmic reticula (Cytopainter). Image analysis of green and red fluorescent pixel intensities (**e**) showed a strong co-localization, particularly at perinuclear regions (**bottom panel**, arrows). Green and red pixel intensities co-occurred (**e**) with 71% of GFP signals co-localized with 82% of red ER signals (*R*^2^ = 0.84) (**Methods)**. **f, g**, GFP fluorescence increases over time at perinuclear ER sites. Single cell analysis of fluorescence intensity over time from perinuclear (**top panel**, *n* = 6) or cytoplasmic (**bottom panel**) regions showed increases over seconds. Fluorescence intensity analysis at perinuclear and cytoplasmic regions of interest in cells expressing PTR showed increasing levels at perinuclear regions compared to within the cytoplasm (sample trace shown in **g**). Scale bars are 30 μm in **a, left panel** and **d,** and **e, top panels,** and 20 μm in **a, right panel** and **e, bottom panel**.

We sought to directly observe protein synthesis at ribosomal sites over time, so we used a live marker for endoplasmic reticulum (ER) in the red channel, Cytopainter. We observed rapid increases in green fluorescence intensity over seconds at perinuclear ER sites which were strongly labeled in red, indicating protein synthesis of the GFP11 at perinuclear ribosomes directly after mRNA export from the nucleus (**Figure 2d, e**). Red and green fluorescence were co-localized throughout the cell (**Figure 2e**), and green fluorescence increased over time only at perinuclear regions compared to the cytoplasm (**Figure 2f, g**). Protein translation of the 26 residues that make up the GFP11 occurs within a few seconds (Ingolia et al., 2011; Karpinets et al., 2006), but even as the protein of interest diffuses away or is translocated into the endoplasmic reticulum during the GFP11 synthesis, the fluorescence event still signifies the ribosomal site of protein translation. Diffusion coefficients for mRNA decorated with ribosomes will vary based on the overall size (Einstein, 1905), but measurements around 0.03 μm^2^ s^-1^ indicate that the ribosome will travel a root mean square displacement of less than 500 nm in 3 seconds (Bakshi et al., 2012; Pichon et al., 2016). Using PTR, there will be a <1 μm spatial variance in detecting the location of the original synthesis event. Traditional methods of protein synthesis reporting using even the fastest folding and maturing fluorescent proteins (which does not account for the time for synthesis of the protein itself) would introduce at least a five minute temporal spread or a 230 μm spatial spread in detection efficiency.

The accumulation of the PTR fluorescence signal over time can eventually make it difficult to detect local protein synthesis events. Possible solutions are to photobleach or to photoconvert the previously generated fluorescent signal. However, photobleaching is undesirable due to the risks of photodamage to the cell and its protein translation complexes, and diffusion of unbleached fluorescent protein into the site. Thus, we split the monomeric photoconvertible fluorescent protein, mDendra2, which normally emits green fluorescence, but can be permanently photoconverted to emit red fluorescence by UV illumination (Chudakov et al., 2007). Mixing purified split Dendra1-10 and Dendra11 peptides *in vitro* produced detectable green fluorescence within seconds (**Figure 3a, left)**, and the fluorescence intensity rose steadily throughout the recording window (**Figure 3a, right)**. UV illumination of the split Dendra1-10 prior to reconstitution did not produce any red fluorescence.

**Figure 3.**
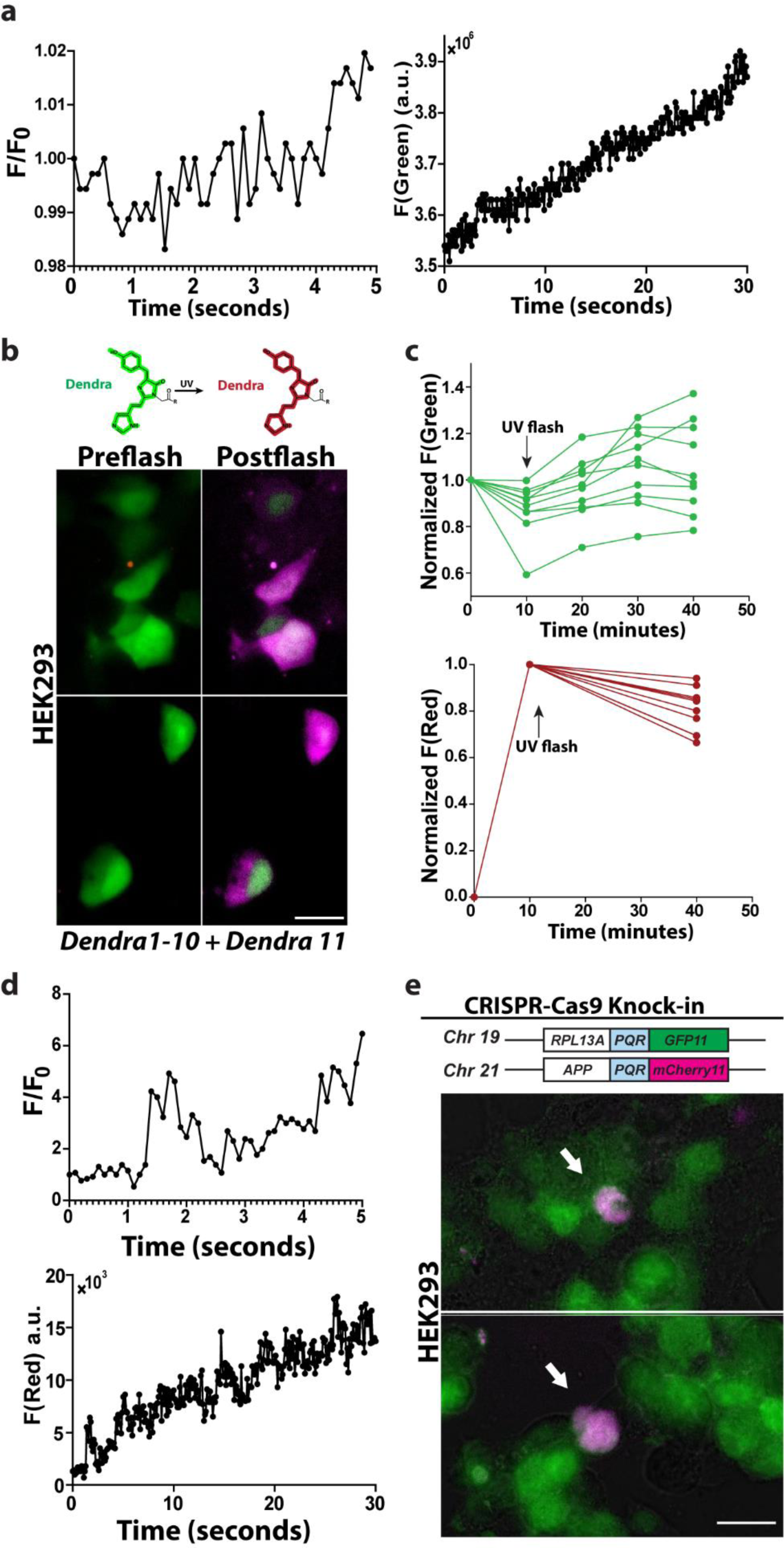
Spectral variants of split PTR. **a***, In vitro* reconstitution of Dendra1-10 and Dendra11 resulted in green fluorescence that was recorded every 0.1 seconds for a duration of 30 seconds. Green fluorescence intensity was detected immediately after mixing of Dendra1-10 lysate and Dendra11 purified peptide at a 200:100 nmolar ratio at room temperature. **b, c**, Split photoconvertible fluorescent proteins can be used with PTR to reset protein synthesis measurements at any time. Reconstituted Dendra2 emits green fluorescence (**b, left panels**), but can be permanently photoconverted to emit red fluorescence upon UV illumination. HEK293 cells co-expressing Dendra1-10 and Dendra11 produce bright green uniform fluorescence that photoconverted to red fluorescence with a 5 second flash of UV light (**b, right panels**). The time course of the green and red fluorescence intensities (**c**) before and after photoconversion (UV flash) shows the relative changes of reconstituted split Dendra2. As protein synthesis continued, Dendra2 green fluorescence increased over time scales of minutes in single cells (individual traces) to values beyond their starting intensity. **d**, Reconstitution of Cherry1-10 lysate and purified Cherry11 peptide resulted in red fluorescence that was detectable immediately after mixing. Fluorescence output was recorded every 0.1 seconds for a duration of 30 seconds. **e**, Multiple genes or alleles can be simultaneously tracked and quantified using other colors of split fluorescent proteins. CRISPR-Cas9 genome editing can insert the small, < 80 bases PTR into genomic loci to measure endogenous protein synthesis. We genome edited HEK293 cells to insert PTRs of GFP11 and mCherry11 into the *Ribosomal Protein L13A* (*RPL13A*) and *Amyloid Precursor Protein* (*APP*) genes, respectively. Green and red fluorescent cells (arrows) simultaneously report the instantaneous protein synthesis of RPL13A and APP over time. Scale bars are 20 μm in **b** and **e**.

By co-expressing these two “Dendra1-10” and “Dendra11” split photoconvertible fluorescent proteins in HEK293 cells, we observed bright green fluorescence (**Figure 3b**). The green fluorescence was then photoconverted by a 5 second flash of UV illumination to red fluorescence (**Figure 3b**). Longer UV illumination to convert more of the existing green fluorescence into red resulted in photobleaching of the red fluorescence signal. Still, new protein synthesis was observed with the increase of green fluorescence over time (**Figure 3c**). Thus, using split photoconvertible fluorescence proteins can enable the precise determination of the moment of protein synthesis, in addition to the rate, by resetting measurement windows at any point in the life of the cell or animal, without the harmful effects associated with photobleaching.

To quantify multiple genes or alleles simultaneously, other split fluorescent proteins (Do and Boxer, 2011; Kamiyama et al., 2016; Kerppola, 2006) can be adapted for PTR for multi-color imaging of protein synthesis. For example, we have previously used CRISPR-Cas9 genome editing to insert *PQRs* with different fluorescent proteins into multiple endogenous genes (Kays and Chen, 2019; Lo et al., 2015) or to track each parental allele (Lo and Chen, 2019). To this end, we used a split red fluorescent protein, mCherry (Fan et al., 2008), to track protein synthesis of two genes simultaneously (**Methods**). Reconstitution of Cherry1-10 and Cherry11 peptides *in vitro* resulted in production of red fluorescence, and this signal increased throughout the duration of the experiment (**Figure 3d)**. The reconstitution of split Cherry occurred immediately upon addition of Cherry11 peptide, but produced an overall weaker signal compared to split GFP and split Dendra2, indicating potentially sub-optimal conditions for the reconstitution reaction.

To track multiple endogenous genes, we used genome editing to insert a *PQR-GFP11* and *PQR-mCherry11* at the end of the coding sequence of the endogenous human *Ribosomal Protein L13A* (*RPL13A*) and *Amyloid Precursor Protein* (*APP*) genes, respectively, in HEK293 suspension cells (**Figure 3e**). The small size of the PTR reporters at 80 bases facilitated efficient integration into the endogenous genes during CRISPR-Cas9 genome editing. These cells were then co-transfected with *GFP1-10* and *mCherry1-10* DNA to co-express the two other split fluorescent protein components. Thus, we were able to simultaneously image endogenous APP protein synthesis in the red channel and RPL13A protein synthesis in the green channel. Ideally however, a universal split1-10 fluorescent protein that alters its emission spectra based on binding of different split11 sequences (Do and Boxer, 2011) would simplify the number of reagents required to perform multi-color PTR.

Localization of mRNA is used as a mechanism for translational regulation. For example, molecular asymmetries can be created by position-dependent translation of mRNAs in oocytes (Nilson and Schupbach, 1999). To verify that PTR can detect protein synthesis of spatially regulated mRNAs *in vivo*, we tracked the production of Gurken protein over time in *Drosophila* oocytes (**Figure 4a**). Synthesis of the Epidermal Growth Factor Receptor ligand Gurken in the anterodorsal corner during the final stages of oocyte maturation specifies the cell fates of only the neighboring follicle cells (Nilson and Schupbach, 1999). We first created transgenic flies that ubiquitously express GFP1-10 from an actin promoter, and flies that express Gurken-PQR-GFP11 under the control of a UAS promoter (**Methods, Figure 4a**). The *Nanos-Gal4* driver was used to express Gurken-PQR-GFP11 in oocytes from females that expressed all three transgenes. We observed rapid increases of reconstituted GFP fluorescence in a restricted anterodorsal region from the nucleus over timescales of minutes in the oocytes (**Figure 4b–d**). These experiments demonstrate that PTR can quantify localized protein synthesis over time.

**Figure 4.**
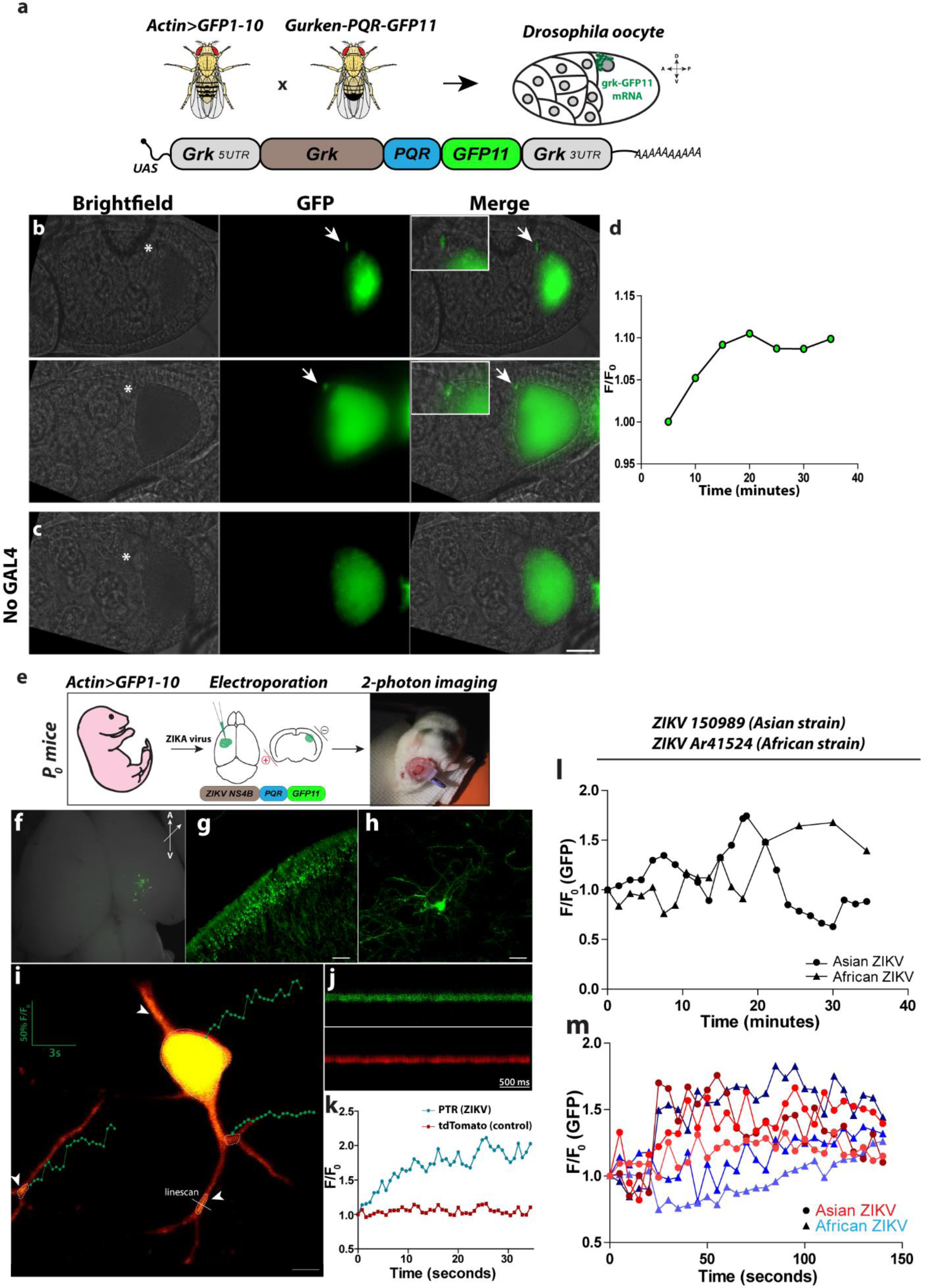
Direct observation of protein synthesis over time *in vivo*. **a**, A transgenic fly line expressing GFP1-10 under the *Actin* promoter (*Actin>GFP1-10*) was created to ubiquitously express GFP1-10 at high levels. We also created *UAS-Grk-PQR-GFP11* flies to visualize translation of local mRNAs. The *grk* transcript contained the native *grk* 5’ and 3’ untranslated regions (UTR) to ensure proper regulation and localization to the anterior dorsal corner of the oocyte near the nucleus, where its translation Gurken initiated. To generate oocytes that express the Gurken-PQR-GFP11, we crossed *nanos-GAL4* to the *UAS-Grk-PQR-GFP11* flies, and then crossed those progeny to the *Actin>GFP1-10* flies. **b**, Representative images of local Gurken translation in *Drosophila* oocytes. Translation of Gurken-PQR-GFP11 produced green fluorescence (arrows) that was always associated within a < 5 μm spread in the anterodorsal region near the nucleus (asterisk) in stage 8 (**upper panels**) and stage 9 (**lower panel**) oocytes (*n* = 6). Oocytes were dissected and imaged over 40 minutes. These results demonstrate the temporal and spatial fidelity of PTR. **c**, Animals expressing the *Actin>GFP1-10* and containing the *UAS-Grk-PQR-GFP11* transgene but without the Gal4 driver is shown to demonstrate the large autofluorescence signal within the nucleus. These negative control animals did not produce the characteristic signal associated with the oocyte nucleus (*n* > 25). Anterior is left, dorsal is up. **d**, Representative example of time-series analysis of GFP fluorescence signals observed at the anterior dorsal corner of the oocyte. GFP intensity increased over timescales of minutes in oocytes. **e,** A transgenic mouse expressing GFP1-10 under the mammalian Actin promoter (Actin>1-10) was generated to constitutively and ubiquitously express GFP1-10 as in **a**. Actin>1-10 pups (P_0_-P_1_) were injected and electroporated with Asian strain *150989* or African strain *Ar41524* Zika virus *ZIKV NS4B-PQR-GFP11* DNA/dye mix (**middle panel**). Animals were allowed to develop for ∼ 14 days and were then imaged using 2-photon microscopy under anesthesia. **f-h,** Green fluorescence was observed 16 hours post electroporation in the injected cortical regions. DNA injected into the lateral ventricles resulted in GFP reconstitution in progenitors and differentiated cells across cortical layers. Green fluorescence was observed in cell somas and projections, indicating high-level expression of GFP1-10 and ZIKV-PQR-GFP11 proteins. Coronal (**g**) and horizontal (**h**) views of postnatal day 1 brain slices are shown. Scale bars are 150 μm in **g**, 30 μm in **h**. **i-k,** 2-photon imaging of local protein synthesis in neurons in the mouse brain *in vivo*. GFP fluorescence increased over seconds at different locations along the neuronal arbor (**i**). Line scans performed at millisecond temporal resolution on different locations along the arbor in the green and red channels (**j**) showed increases in GFP fluorescence but not the tdTomato control plasmid (**k**) (representative scanline and scan shown). Arrowheads indicate regions along the arbor where increased Zika virus NS4B production was observed. Dashed lines delineate the regions of interest used to generate the overlaid traces. Scale bar is 10 μm in **i**. **l, m,** The subcellular protein translation dynamics of African (strain Ar51524) and Asian (strain 150989) ZIKV proteins can be tracked over varying timescales (representative traces from different dendrites over minutes and seconds in **l** and **m**, respectively) in the living animal.

Next, we sought to track and quantify local protein synthesis in neurons *in vivo*. We created a transgenic mouse that constitutively produces high levels of GFP1-10 protein driven by the *actin* promoter in all tissues throughout the life of the animal (**Methods**). We then used PTR with GFP11 to examine the protein synthesis dynamics of two strains of the Zika virus (*ZIKAV*) protein NS4B in the perinatal mouse brain. The Zika virus is an RNA virus that can cause Zika fever in humans, and fetal microcephaly in pregnant humans (Brasil et al., 2016). We used *in vivo* electroporation in neonatal (Postnatal day P0-P1) mice (**Figure 4e**) to express the NS4B Zika virus protein with PQR-GFP11 flanked by the native 5’ and 3’ untranslated regions, *5’UTR-NS4B-PQR-GFP11-3’UTR*, from either the African Ar41524 or Asian 15098 strain. Co-electroporation of a DNA plasmid of the red fluorescence protein tdTomato was used as a cellular marker and its production dynamics to compare traditional and PTR-based reporters. We observed green fluorescence in cell somas and neurites as early as 16 hours post electroporation (**Figure 4f–h**). This indicated that GFP11 peptides were co-produced with the viral NS4B protein and that sufficient GFP1-10 protein was being produced under the Actin promoter in the transgenic mice. Using 2-photon laser scanning microscopy, we tracked the *in vivo* production of NS4B protein in neurons by measuring changes in green fluorescence over time (**Figure 4i–k, Supplementary Movie 1**). Images were acquired at varying time intervals (400 milliseconds to 5 minutes) to verify that the time course of fluorescence signals were not due to changes in animal movement, position, photobleaching or imaging depth (**Figure 4k–m**). We observed bursts of NS4B protein synthesis over seconds to minutes, and different subcellular locations. For example, local increases of fluorescence intensity were found at perinuclear regions and along proximal dendrites (arrowheads in **Figure 4i**). We also observed localized increases in protein synthesis within dendrites, and line scan imaging across these spots showed increases in GFP fluorescence over milliseconds (**Figure 4j**). Importantly, over these timescales we observed no significant changes in tdTomato red fluorescence, which demonstrates the temporal and spatial sensitivity of PTR over traditional full-length reporters *in vivo* (**Figure 4k**).

Long maturation times of traditional fluorescent proteins result in a broad spread of fluorescence observed in a soma (**Figure 4k**) due to their diffusion from the sites of translation during fluorescent protein maturation. Over-expression of PTR plasmids will cause a continual increase in fluorescence during the first few days of transfection (**Figure 1c, d**), but local protein synthesis (whether by random or trafficked mRNA distribution) is still detectable (**Figures 2d, 4k**). Using PQR to track endogenous protein synthesis by genome editing a PQR into an endogenous gene has shown that the PQR fluorescence does not always constantly increase (Lo and Chen, 2019).

## Discussion

Protein synthesis of a PQR-GFP11 reporter requires approximately 2 seconds (6 amino acids/second) (Kramer et al., 2009; Ross and Orlowski, 1982). In the presence of GFP1-10, the split proteins reconstitute in milliseconds to emit a quantitative green fluorescence signal to indicate the time and location of protein translation within the cell. Synthesis of the GFP11 peptide requires ∼ 3 seconds from the moment of initiation of GFP11 translation and the reconstitution and emission of fluorescence. Whether or not the upstream protein diffuses away from the site of translation, translation of GFP11 will mark the original site of mRNA translation, unless the RNA-bound ribosome diffuses away. Polysomes bound to mRNA diffuses at 0.03 μm^2^ / second (Bakshi et al., 2012; Pichon et al., 2016), and thus the 3 second delay in detecting the initial protein translation event will produce a spatial error of ∼ 350 nm. Using genome editing to insert the PQR-XFP11 allows for endogenous tracking of protein synthesis, but may alter mRNA localization and degradation. However, it is unlikely to alter the translation speed (i.e., codon recognition efficiency) of the target gene of interest. Similar to the PQR technique, insertion of the PQR-XFP11 downstream of the gene of interest will leave 21 amino acids on the C-terminus of the target protein, unless the XFP11-PQR is inserted upstream of the target gene (Lo et al., 2015). Another disadvantage to examining endogenous protein expression is that both components are required, the PQR-XFP11 and the XFP1-10 animal, such as the transgenic flies or mice that we generated (**Figure 4**).

Our results suggest that the kinetics and efficiency of the reconstitution reaction can be improved, as different variants of the GFP11, Dendra11, and Cherry11 peptides produced different rates of reconstitution. Typical protein-protein reactions such as antibody-protein associations contend with rotational and relative translation (shifting between atomic interfaces) constraints and have association rate constants, k_on_, around 10^5^ M^-1^ s^-1^. Given that XFP1-10 and XFP11 are split from the same protein and the XFP11 is a small peptide, it is likely that their association rate constants are limited by diffusion and less by geometric constraints. Their association does not require movement of protein domains and their interfaces may be oriented with favorable electrostatic interactions such as complementary charges distributed between the two (Schreiber et al., 2009). Minor differences in the solubility, charge and size of the XFP11 peptide can affect the efficiency of reconstitution, and ultimately the properties of the reconstituted protein. Therefore, screening for XFP11 peptides that result in the most sensitive reconstitution will certainly improve PTR in the context of monitoring local protein translation events. Split fluorescent protein components that fail to reconstitute, or reconstitute but fail to fluoresce can theoretically affect the spatial, temporal and quantitative accuracy of the PTR reporters. While we did not observe such problems in our experiments, their occurrence is difficult to predict or estimate *in vivo*. In addition, our measurements are of ensemble kinetics, and thus, stochastic single-molecule behaviors may apply at extremely high resolutions or at very low concentrations. However, the finding that different variants of green, red, and photoconvertible XFP11s reconstitute differently suggests that undiscovered split fluorescent protein variants will outperform currently available ones.

Our results using PTR to examine differences in local protein synthesis in the living animal demonstrate its applicability as an infectious disease model in addition to examining dendritic protein synthesis. Multi-color subcellular imaging of local protein synthesis can be useful in examining pathogen-host interactions, related genes and pathways, or different parental alleles. For example, tagging of a viral genome with PTR and subsequent infection of a GFP1-10 transgenic mouse with a CRISPR-Cas9 knockin of *IFIT2-PQR-RFP* (*IFIT2* is an interferon responsive gene) allows for real-time quantification of the viral infection spread in single neurons with GFP (e.g., **Figure 4**), while also observing the mouse’s immune response (*IFIT-PQR-RFP*) in the red channel. In another example, multi-color imaging with CRISPR-Cas9 knock-in of PTRs into the maternal and paternal allele (Lo and Chen, 2019) will allow real-time imaging of each allele’s protein product. Thus, traits or disorders that are associated with a specific allele (e.g., an autism-associated mutation in the glutamate receptor subunit GluA1) can be imaged throughout a single neuron to reveal molecular asymmetries, or cellular heterogeneities globally across an organ in the living animal.

## Acknowledgments

The authors thank Chiu-An Lo and Farida Emran for assistance with experiments and analysis, and Laura Nilson for assistance with Gurken experiments. This work was supported by grants (to B.E.C.) from the Natural Sciences and Engineering Research Council of Canada, and the Canadian Institutes of Health Research (148882).

## Author Contributions

B.E.C. designed the experiments and supervised the project. I.K. performed experiments and analyzed the data. I.K. and B.E.C. wrote the manuscript.

## Competing Interest Statement

B.E.C. and I.K. are inventors on a patent application describing the system and materials in this manuscript.

The data that support the findings of this study are available from the corresponding author upon request. Correspondence and requests for materials should be addressed to

**Supplementary Information** is provided as three supplementary movies and seven supplementary figures.

## Methods

### Experimental Design

We sought to design split fluorescent proteins that could reconstitute rapidly, with one split portion able to be synthesized rapidly and the other remain non-fluorescent until reconstitution. The biological experiments were designed to validate the PTR system and demonstrate its utility in different model systems. All experiments were performed in experimental quadruplicates at minimum. Biological/experimental replicates were defined as all cells within a plate *in vitro*, or single cells within an animal *in vivo*. Multiple regions of interest (ROIs) within and across cells in a plate were averaged together for *in vitro* experiments, and multiple ROIs were averaged per single cell in *in vivo* experiments. No outlier data points were encountered and no data were chosen for exclusion. All cells, animals, fields of view, and groups used in this work were selected for analysis at random. The investigators were blind to genotypes and conditions where permitting, and all data analysis was performed with investigators blind to experimental parameters.

### Split Fluorescent Reporter DNA constructs

Residues 1-213 of GFP, corresponding to the first 10 beta strands of GFP (GFP1-10), were amplified and cloned from evolved superfolderGFP. GFP11 was generated by amplifying and cloning the last 15 residues of GFP into pCAG. To stoichiometrically co-express GFP1-10 or GFP11 with other proteins of interest, a PQR construct was added in-frame upstream or downstream of the GFP1-10 or GFP 11 sequence depending on the desired orientation. For extracellular membrane-bound expression of GFP1-10, the complete Neuroligin-1 signal sequence, in addition to portions of the Neuroligin-1 extracellular, transmembrane, and intracellular anchoring domains were fused to the N-terminus of GFP1-10 (Landgraf et al., 2012). For electrophysiology experiments, a PQR-GFP11 reporter was placed downstream of the ShakerRFP coding sequence to generate ShakerRFP-PQR-GFP11, and cloned into pCAG. Split mCherry was generated as described (Fan et al., 2008). Split Dendra2 was generated by separating mDendra2 (Evrogen, Moscow, Russia) at Proline 191 to generate Dendra1-10. The remaining 39 residues (Dendra11) were cloned into pCAG or placed downstream of a PQR to generate a Dendra11 protein translation reporter. Split mCherry was generated according to Fan et al., (Fan et al., 2008). For ZIKV experiments, the portion corresponding to full length NS4B was cloned from African ZIKV strain *Aedes africanus*/SEN/DakAr41524/1984 (Genbank: KX601166.2) or Asian ZIKV strain isolate 15098 (Genbank: MF073359.1), upstream of PQRGFP11 into pCAG. The control plasmid tdTomato was expressed using pCAG. For transgenic animals, the sequence encoding GFP1-10 was cloned downstream of the mouse or fly *ActinB* promoter. For mouse transgenesis, *Actin>GFP1-10* was cloned into pCAG and injected into mouse blastocysts according to standard transgenic practices (McGill Core Transgenic Facility). Similarly, fly *Actin>GFP1-10* was cloned into a modified version of pCFD3 (Addgene 49410) and injected into fly embryos (BestGene, Inc) (**Supplementary Figure 2**). If GFP1-10 is expressed at even 1% of the levels of Actin, which is only 10 times the median copy number for a protein in a mammalian cell (Schwanhausser et al., 2011), there still will be > 500 GFP1-10 molecules per 1 fL within the cell for GFP11 to bind.

### XFP1-10 protein production and extraction

DNA encoding XFP1-10 protein was transformed and expressed under the control of an arabinose inducible promoter in Escherichia coli strain BL21(DE3) (New England BioLabs, Ipswich, MA). Cells were grown in Luria broth medium to an initial optical density of 0.2, at which point induction of protein production was initiated with 0.2% L-Arabinose and cells were further grown at 37°C for an additional 16 hours with shaking at 225 rpm to encourage inclusion body formation. Cultures were harvested using centrifugation and XFP1-10 was purified from inclusion bodies by resuspension with TNG buffer (100 mM Tris-HCl (pH 7.4), 150 mM NaCl,10% glycerol vol/vol) containing 0.5 mg/ml lysozyme, 50 units of DNase I. The lysate was then incubated at 37°C for 25 min. Crude lysates containing XFP1-10-rich inclusion bodies were separated using centrifugation at 16,000g at 4°C. Inclusion bodies were lysed using B-Per (Thermo-Fisher) and sonication, and XFP1-10 protein was collected and filtered using a 0.22 μm filter before concentration with 10,000 molecular weight exclusion columns. We used this lysate as concentrated XFP1-10, but because this was not pure XFP1-10, this would underestimate our k_obs_ measurements.

### XFP11 Peptides

Variants of the XFP11 peptide were chemically synthesized with >75% purity (Genscript, Piscataway, NJ). The amino acid sequences of the GFP11 peptides were: GFP11v1: RDHMVLHEYVNAAGIT, GFP11v2: RDHMVLLEFVTAAGIT and GFP11v3: RDHMVLHEFVTAAGIT (see Table 1 for full sequence list). Lyophilized peptides were resuspended in water to >10 mg/mL and frozen at -20°C. For extracellular membrane GFP reconstitution in HEK293 cells, GFP11 peptide was dissolved into the culture medium at a final concentration of 50 μM and cells were returned to a 37°C incubator for two hours before live imaging.

### *In Vitro* Protein Reconstitution

*In vitro* fluorescence complementation was performed by mixing purified XFP1-10 protein and chemically synthesized XFP11 peptides and the fluorescence intensity of the reaction was collected with a StepOnePlus real-time thermal cycler (Thermo-Fisher). 3 mM XFP1-10 in TNG buffer or PBS (varied pH) was added to wells of a 96-well microplate coated with 1 mM bovine serum albumin and allowed to equilibrate for 60 seconds. XFP11 peptide was added according to different final peptide concentrations and the microplate was immediately loaded into the fluorescence reader. The fluorescence intensity was measured every 10 seconds for 45 minutes at 32°C or 37°C with 495 nm excitation and 520 nm emission. The fluorescence intensity was normalized to the initial fluorescence intensity to express relative fluorescence increase upon fluorescence reconstitution. We verified that split GFP, mCherry, and Dendra2 had similar rapid kinetics, and that UV illumination did not photoconvert the split Dendra1-10. We illuminated split Dendra1-10 with UV excitation for 30 seconds to one minute and then performed complementation with split Dendra11 in multiple experiments and did not observe any red fluorescence. Different XFP11s do not cross-react with orthogonal XFP1-10s, although this is a future goal (Do and Boxer, 2011).

Standard curves for XFP fluorescence measurements were generated by either using reconstituted XFP or XFP purified from *E.coli* using GFP specific chromatography columns (Bio-Rad). XFP protein concentration was determined using the Bradford assay and absorbance readings at 280nm with a NanoDrop 2000 (Thermo-Fisher). Samples were serially diluted (1:10 or 1:5) and 10 μL samples were imaged to reduce any non-linear fluorescence excitation effects. For sub-second kinetic measurements, a Fluorolog-3 spectrophotometer (Horiba) fitted with a stopped-flow dispenser was used to simultaneously dispense and mix known volumes of XFP1-10 and XFP11 purified proteins at room temperature. Total fluorescence intensity (in photon counts / time) was collected every 0.1 or 0.01 seconds for varying durations. In the Figures, fluorescence normalized to initial fluorescence (F / F_0_) is used on *y*-axes to show initial rapid changes in fluorescence; the raw fluorescence intensity values in arbitrary units (a.u.) is used on *y*-axes to show the raw changes across the overall time. We also used standard curves of known nM concentrations of reconstituted split GFP (**Figure 1h, top left**) to verify that a standard fluorescence microscope can detect PTR signals over a range of physiological concentrations.

The median copy number of proteins in mammalian cells is 50,000 (Schwanhausser et al., 2011), which is 80 nM for an average 1000 fL cell. The rates of various XFP1-10 and XFP11 binding reactions are dependent on the concentration of XFP11, the concentration of XFP1-10, the association rate constant, *k_on_*, and the dissociation rate constant, *k_off_*. Thus, it is important to take all of these factors into account when determining the speed of PTR. The equation used for calculating fluorescence over time (*t*) based on k_obs_ was F(*t*) = F_max_ × (1-1/*e^kt^*) + F_auto_, where F_max_ is the maximum fluorescence measured, *k* is k_obs_, and F_auto_ is the autofluorescence from unbound XFP1-10 or any other sources. Autofluorescence values for the XFP1-10 fluorophores were only detectable when compared to a blank (**Figure 1f**). Cells transfected with XFP1-10 alone were not noticeably brighter than non-transfected cells (**Supplementary Figure 3**).

### Cell Culture

HEK293 cells were cultured at 37°C under 5% CO_2_ in Dulbecco’s Modified Eagle Medium, supplemented with 10% fetal bovine serum (Wisent), or for human NPCs, or mouse cortical neurons, in NPC Medium or neural differentiation Medium (Sigma-Aldrich). Media were supplemented with 100 units/mL penicillin (Thermo-Fisher) and 100 μg/mL streptomycin (Thermo-Fisher). Mammalian cells were transfected with 3.5 μg of plasmid DNA in 35 mm dishes using Lipofectamine 3000 (Thermo-Fisher). Cells were imaged 24–36 hours later. For extracellular GFP fluorescence reconstitution, HEK293 cells were transfected with constructs expressing GFP1-10 tagged to the transmembrane and extracellular domains of the cell surface molecule Neuroligin-1 and incubated for 24–36 hours. Cells displaying GFP1-10 on the extracellular side of the cell membrane were incubated in culture medium containing 50 μM GFP11 peptide for 3 hours at 37°C before live imaging. GFP1-10 protein was allowed to accumulate for 24 hours before transfection of GFP11 constructs. For genome editing experiments, 800 ng of CRISPR-Cas9 plasmid DNA were co-transfected with 800 ng of single stranded oligonucleotide repair templates in 12-well plates. After 2–7 days, cells were non-enzymatically dissociated and seeded on glass coverslips and prepared for imaging and electrophysiology experiments.

To verify that the PTR fluorescence signals were due to protein synthesis, we blocked protein synthesis using Cycloheximide to bind ribosomes. Cycloheximide was purchased from Sigma–Aldrich (C4859-1mL, 100mg / mL) and dissolved in ethanol to a stock concentration of 20 mg / mL. Cycloheximide was added at a final concentration of 1–10 μg / mL in the cell culture media, 5 minutes prior to imaging. Protein synthesis inhibition in HEK293 cells using Cycloheximide occurs rapidly within minutes of addition to the cell culture media (Kearse et al., 2019). Addition of Cycloheximide decreased PTR fluorescence within minutes, compared to standard fluorescent protein transfections, or no Cycloheximide controls (**Supplementary Figure 4**).

### Endoplasmic reticulum and ribosome staining

To visualize endoplasmic reticula (ER), HEK293 cells were transfected with split GFP PQR reporter constructs and stained (live or fixed) with the ER-specific stain Cytopainter (Abcam). Stained cells were imaged in the green and red channels to examine the co-localization of reconstituted GFP and red ER signals. Colocalization of green and red signals was determined by calculating the Pearson’s and Mander’s correlation coefficients for overlapping green and red pixel intensities. Individual z-planes were background subtracted and thresholded to remove the lowest and highest pixel intensities. Ten regions of interest (ROIs) within the cell and excluding background and nuclear regions were used for analysis. Both Pearson and Mander’s colocolization coefficients were independently obtained and cross validated using Coloc2 (ImageJ) and BioImageXD (Kankaanpaa et al., 2012). Average (with standard deviation) GFP fluorescence time courses at perinuclear ribosome sites versus cytoplasmic sites are shown in **Supplementary Figure 5**.

### Image Acquisition

All imaging experiments were performed in experimental quadruplicates at minimum. Fluorescence and brightfield microscopy were performed using a Zeiss AxioScope A1. All images were acquired at 1388 x 1040 pixels using a 40× water objective, N.A. 1.0 (epifluorescence). Fluorescence emission was detected using a charge-coupled device (CCD) camera (MRm). All image acquisition parameters were fixed for each imaging channel for exposure time, excitation intensity and gain. Cells that were dimmer or brighter than the fixed initial acquisition dynamic range were not included for analysis. Multiple focal planes were imaged for maximal *z*-projection analysis. Time-series images were collected using an open-shutter video configuration in ZenLite (Zeiss). Images were acquired every 167 milliseconds with exposure times of 260 milliseconds. For *in vivo* 2 photon imaging, juvenile mice (P_10_-P_20_) were anaesthetized using ketamine/xylazine/acepromazine and a 4x4 mm square window was made through the skull using a dental drill fitted with a 0.45mm drill burr. The meninges were carefully removed and the exposed brain was covered with 1% optically clear agar and sealed with a No.1 round coverslip. The coverslip was secured to the skull using dental cement and a mounting fixture was also placed (**Figure 4e, right panel**) to facilitate securing the mouse under the objective to minimize animal movement, breathing artifacts and vibration. Images were acquired using a custom-built 2-photon laser scanning microscope at 512 x 512 pixels using a 40× water objective, N.A. 1.0.

### Image Analysis

Image analyses were performed blind with respect to condition or genotype. Images were selected for analysis based on identification of healthy cells and low background. Fluorescence pixel intensities were measured in several random ROIs within the target cellular region using a custom written program in MatLab (MathWorks) or ImageJ. Average pixel intensities were calculated from five ROIs of 7x7 pixels for measurements within the cytoplasm, perinucleus, and nucleus, and 3x3 pixels for measurements within the plasma membrane (e.g., ShakerRFP). All signal intensities were background subtracted from the average of three 10x10 ROIs immediately surrounding the cell. For time-series image analysis, background was considered as the region immediately adjacent (<15 μm) to the perinucleus, or cytoplasm. Perinucleus was defined as a 5 μm ring around the nucleus, and cytoplasm was defined as any region beyond that ring without ER. Each time point from each experiment consisted of at least seven ROIs from at least four cells. Data points used for graphs were the average of the experimental replicates (minimally in quadruplicate).

For co-localization analysis, Mander and Pearson’s correlation coefficients were calculated for individual z-slices. Using the ribosome marker in the red channel with PTR GFP experiments, green and red pixel intensities were confirmed to colocalize, with 71% of above-threshold GFP signals co-localized with 82% of above-threshold red signals. Thresholded Mander’s (tM) coefficients tM1 = 0.71, tM2 = 0.82, Pearson’s *R^2^* = 0.84, and Costes p-value = 1 (*i = 100, n = 10 ROIs*).

Kinetic increases in fluorescence from timelapse or video data were plotted was Ft/F_0_. For ZIKV translation dynamics experiments, autocorrelation analysis of the single cell fluorescence time courses was used to provide an unbiased estimate of the temporal variation in protein translation events between ZIKV strains. We sought to examine differences in protein synthesis rates between two strains of ZIKV, the ancestral African strain (Ar41524) and the divergent Asian lineage strain (15098). We hypothesized that differences in the 5’ and 3’ mRNA UTRs or the nonstructural NS4B portion (involved in virus replication and evasion of host innate immunity) that is codon usage divergent between the two strains, would alter local protein synthesis dynamics in mammalian neurons. We performed an autocorrelation analysis on the fluorescence time course as an unbiased estimate of burst duration using different regions of interests within the soma and dendrites (30 minute duration sample, *n* = 3) (**Supplementary Figure 6**). A single exponential decay was fit to the autocorrelation function and resulted in an average time constant of τ = 8.84 ± 9.48 seconds for Asian ZIKV *150989* (*R^2^=0.99, n=3 cells, 2 animals*) compared to τ = 12.86 ± 11.23 seconds for African ZIKV *Ar41524* (*R^2^=0.97, n=4 cells, 2 animals*) (mean ± SD, *p* = 0.54, *n* = 7 cells) (**Supplementary Figure 6**). Thus, we found no significant difference in NS4B protein production between the ancestral versus divergent ZIKV strains in neurons. Individual green and red channel images for **Figures 2** and **3** are shown in **Supplementary Figure 7**.

### Electrophysiology

Standard whole cell voltage clamp was used to record potassium currents from HEK293 cells _6_. Cells were maintained at 25°C in extracellular solution containing 140 mM NaCl, 10 mM CaCl2, 7.5 mM KCl, 10 mM HEPES, and 10 mM glucose at pH 7.4, 319 mOsm during recordings. Patch electrodes were pulled from standard wall borosilicate glass (BF150-86-10, Sutter instruments) with 3–5 MΩ resistances. The intracellular pipette solution was 120 mM KCl, 2 mM MgCl2, 1 mM CaCl2, 2 mM EGTA, 20 mM HEPES, and 20 mM sucrose at pH 7.23, 326 mOsm. Whole cell currents were low pass filtered at 10 kHz and measured using an Axopatch 200B amplifier (Axon instruments) and recorded using a DigiData 1200 with pClamp9 software (Molecular Devices). Cells were held at -80 mV and then given +20 mV steps of 45 ms. To accurately compare I-V curves and current data across cells and experiments, the steady-state current was divided by the membrane capacitance (mean Cm = 15 pF, *n* = 3), and current density (pA/pF) was used for comparisons. Consistent cell capacitance, and membrane and access resistances were verified before and after recordings.

### Genome Editing using CRISPR-Cas9

Guide RNAs were designed as 20 bp DNA oligonucleotides and cloned into *pX330* (Addgene 42230), and co-transfected with a circular PQR repair template using Lipofectamine LTX (Life Technologies). All CRISPR-Cas9 guide RNAs were tested for activity using SURVEYOR Nuclease and SURVEYOR Enhancer S (Transgenomics) on extracted genomic DNA. Re-annealed products were analyzed on 4%–20% Novex TBE polyacrylamide gels (Life Technologies). To construct the repair templates, the gene sequences were identified in the human genome using GeneDig (https://genedig.org, (Suciu et al., 2015) and then PCR amplified. Repair templates were constructed by placing *PQR-XFP* between homology arms specific for the genes. The homology arms lacked the promoter, which prevented expression of the *PQR-XFP* until in-frame genomic integration within an active coding gene. Left and right homology arms were each one kilobase long. Cellular fluorescence from PQRs was observed four days post-transfection.

### *Drosophila* oocyte imaging

We generated a transgenic fly (BestGene, Inc) containing *UAS-Grk5’UTR-Grk-PQR-GFP11-Grk3’UTR*, which included the RNA transport and localization elements in the 5’ and 3’ untranslated regions (UTRs) to ensure proper translational regulation (Saunders and Cohen, 1999). These flies were crossed to *nanos-GAL4* and *Actin>GFP1-10* flies to create the triple transgenic flies with *nanos-Gal4* driving the *UAS-Grk-PQR-GFP11* in a background of *Actin>GFP1-10*. This ensured the maternal expression of these mRNAs within oocytes. *Drosophila* ovaries were extracted in Schneider’s medium to reveal the egg chambers containing the various stages of development. Stage 8 – 10 oocytes were selected for fluorescence imaging and showed a crescent shaped green signal that always localized to the corner near the nucleus (*n* = 6).

### Neonatal Brain Electroporation

All animal experiments were performed in accordance with the institutional guidelines. P_0_-P_1_ *Actin>GFP1-10* mouse pups were anesthetized on ice until no reflexes were observed. 2 μg / μL *NS4B-PQR-GFP11* DNA (from Asian ZIKV strain *150989* or African ZIKV strain *Ar41524*) and *tdTomato* DNA dissolved in water was mixed with 0.1% FastGreen dye and loaded into a 1.5 / 0.786 mm (O.D. / I.D.) pulled and beveled borosilicate glass micropipette (Sutter Instruments). The micropipette was inserted into the right lateral ventricle or ∼1.5 mm into the right cortical hemisphere, ∼ 1.5 mm laterally from bregma (**Figure 4e**). Around 200 nL of DNA and dye solution was injected and allowed to diffuse for three minutes. Platinum tweezer electrodes (Nepagene) were then placed around the pup head such that the negative electrode contacted the injected side of the head. A drop of PBS was used under the electrodes to transmit the pulses. Two sets of nine pulses (100 V, 50 ms, 950 ms apart) separated by three seconds were delivered, and the animal was allowed to recover on a 37°C heated blanket. The pups were returned to their mother once awake and active.

### Statistical Analysis

Pearson’s linear correlations were calculated by fitting the data to a simple linear regression model, with the coefficient of determination, *R^2^*. Kinetic reconstitution traces were fit with a simple one-site binding model with the coefficient of determination R2 using Prism (GraphPad). We used the *F* test to test the null hypothesis that the variables were independent of each other and that the true *R^2^* value was 0 for both linear and nonlinear models. Autocorrelation analysis was performed using custom-written programs in MatLab (MathWorks, Natick, MA). The autocorrelation function was fit to single-phase exponential decay models using Prism (GraphPad). A Mann-Whitney *U* test was used to test the null hypothesis that protein synthesis burst rates between the two Zika virus strains are not different.

## Supplementary Information

**Supplementary Figure 1.**
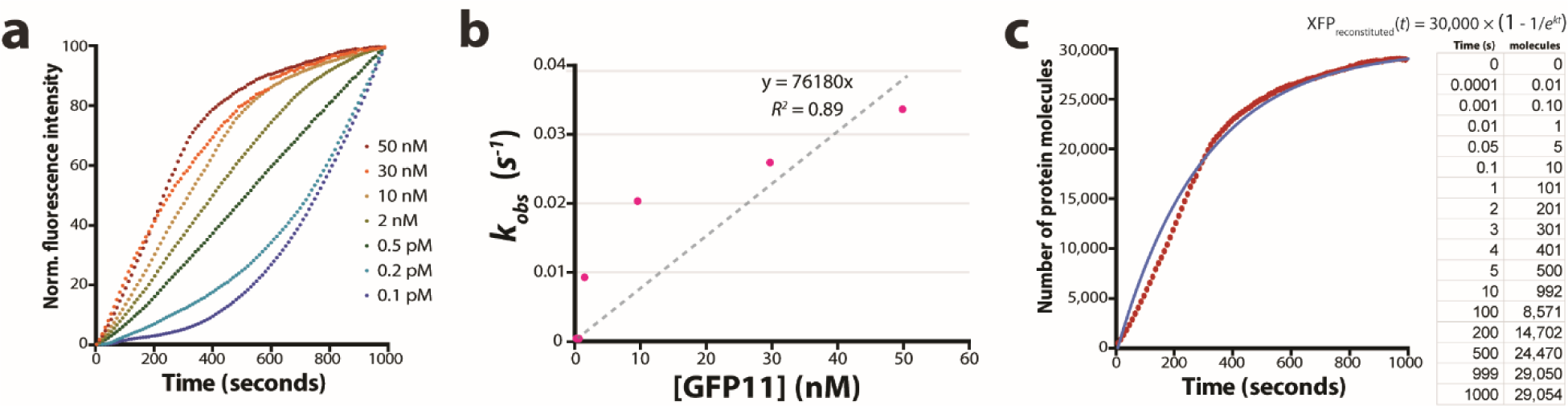
SplitGFP reconstitution occurs rapidly. **a**, Reconstitution of GFP11 with excess GFP1-10 occurs extremely rapidly. Varying concentrations of GFP11 peptide were mixed with 3 mM GFP1-10 and fit with a one phase association curve (same data as in **Figure 1g**) to determine k_observed_ for each concentration of GFP11. The k_obs_ for 50 nM of GFP11 is 0.003 s^-1^ with a half-time of 210 seconds. Thus, at 50 nM protein, which is 40% less than the median copy number of proteins in a mammalian cell, in the first second of reaction time more than 100 molecules of GFP11 have bound to GFP1-10. **b,** The observed rate constant (k_obs_) for GFP11 binding to GFP1-10 is a function of GFP11 concentration. Each GFP11 concentration curve in **a** was modeled as a pseudo-first order reaction and fit to a one phase association curve to calculate k_obs_ (*R^2^* > 0.98, *p* < 0.001, for each concentration, **Methods**). Plotting k_obs_ versus [GFP11] with a linear fit gives the equation y = 761803x, with the slope equal to k_on_ and a y-intercept of zero. Thus, the association rate constant (k_on_) is 7.6 × 10^5^ M^−1^ s^−1^. The K_d_ was 481 ± 116 pM, meaning a k_off_ of 3.7 × 10^-7^ s^-1^. We did not experimentally measure k_off_, but others have found that it is in the many hundreds of hours (Do and Boxer, 2011). The speed or rate at which the XFP11 binding to XFP1-10 reaction occurs *in vitro* or *in vivo* is not the same thing as the observed rate constant, k_obs_, which is also not the same thing as the association rate constant, k_on_. The observed rate constants, k_obs_, in **a** are dependent on the XFP11 concentrations, and the association, k_on_, and dissociation, k_off_, rate constants. The association and dissociation rate constants are a property of the XFP11 and XFP1-10 molecules and are fixed for each pair of XFP constructs, which we measured by plotting multiple k_obs_. The association rate constant, k_on_, of 10^5^ M^−1^ s^−1^ indicates high affinity and rapid binding, or diffusion-limited reactions, but it is important to note that association rate constants are not slow or fast, they are simply small or large. Instead, the speed of various XFP11 and XFP1-10 binding reactions are dependent on the concentration of XFP11, the concentration of XFP1-10, the association rate constant, k_on_, and the dissociation rate constant, k_off_. Thus, it is important to take all of these factors into account when determining the speed, or rate, of PTR. **c,** The speed of XFP11 and XFP1-10 binding changes over time *in vitro* as reactants are used up. The speed of XFP11 and XFP1-10 binding is dependent on the concentration of XFP11 and the concentration of XFP1-10 (and the association and dissociation rate constants, k_on_ and k_off_, respectively). The reconstitution of XFP11 and XFP1-10 is plotted with the total number of molecules shown in the ordinate and time on the abscissa. The formula for the exponential curve fit to the data is shown and examples of number of molecules reconstituted over time in the table inset. Thus, the speed of reconstitution can change over time as the concentrations of XFP11 and XFP1-10 change, so, using the formula or table we see that in the first 100 milliseconds 10 molecules of XFP11 and XFP1-10 have reconstituted but in the 1 second between 999 and 1,000 seconds, only 4 molecules have reconstituted. The time resolution of PTR binding detection will change as XFP11 or XFP1-10 concentration changes over time. For a physiological perspective, 1 molecule in an average 1000 fL cell (1 divided by Avogadro’s constant divided by 1000 fL) is 1.7 pM. 10,000 molecules in an average cell is 17 nM. We measured k_obs_ for XFP11 concentrations far lower than 1 molecule in an average cell, and far lower than an expected K_d_. The median copy number of RNA molecules in mammalian cells is 17, with a narrow distribution of RNA copy numbers for different genes (Schwanhausser et al., 2011). Proteins are about 3,000 times more abundant than RNA, with the median copy number of proteins at 50,000, which is 80 nM (Schwanhausser et al., 2011). The range of RNA concentration in a cell is between 1 pM to 1 nM, but protein concentrations are between 1 nM to hundreds of µM. The median half-life for RNA is 9 hours, and thus each RNA strand must synthesize thousands of proteins in its lifetime. The rate constant observed in experiments, k_obs_, is dependent on ligand concentration, and so a large k_obs_ can be measured by changing the observation volume and focusing on local protein synthesis. Large local concentrations of protein can occur, for example, in dendrites or at the nuclear membrane.

**Supplementary Figure 2.**
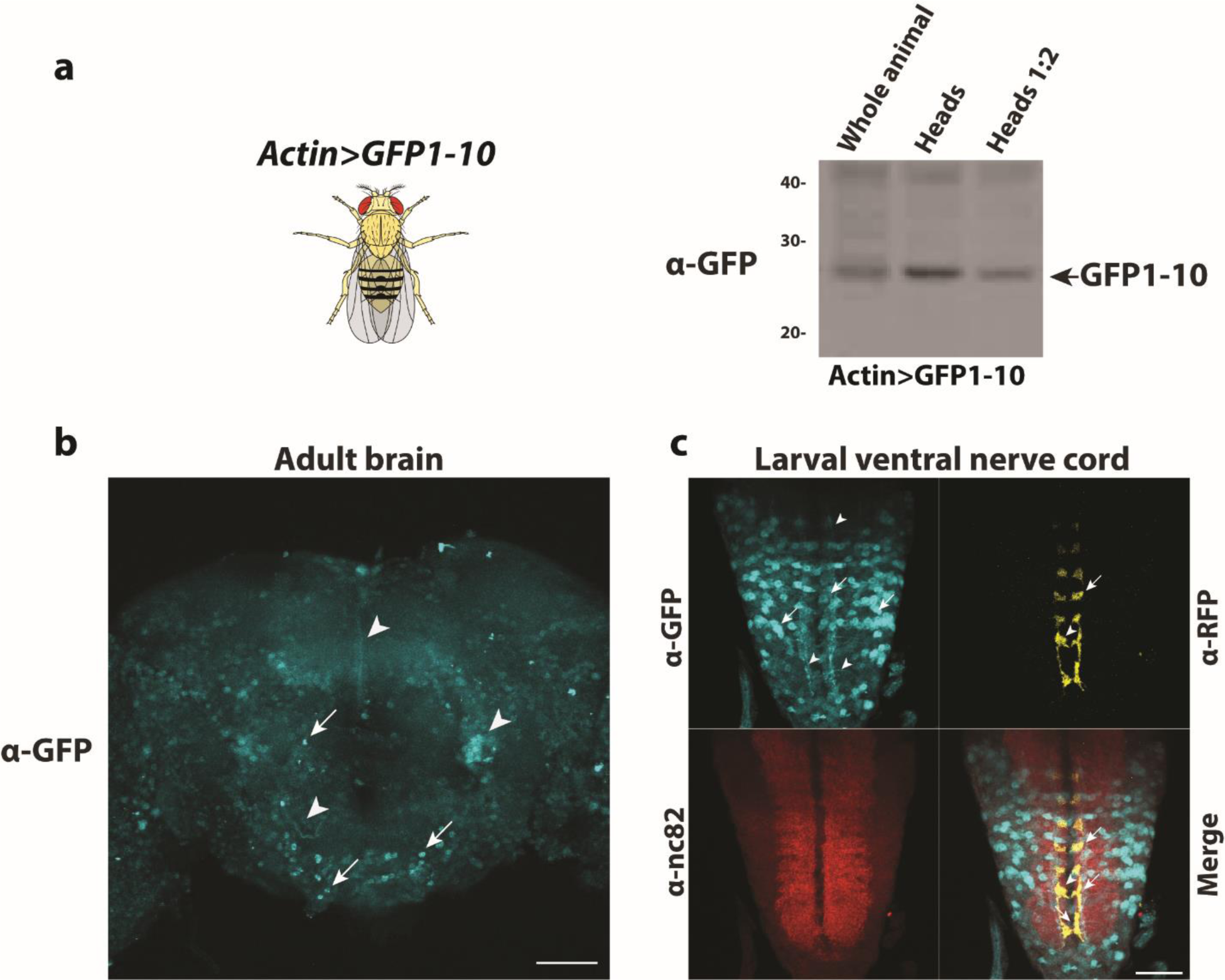
GFP1-10 is expressed at high levels in transgenic animals. **a**, Immunoblot performed using α-GFP antibody on whole fly and head extracts. The 24 kDa predicted GFP1-10 protein is expressed in flies and runs at the expected size. **b-c** Immunohistochemistry showing ubiquitous and high-level expression of GFP1-10 protein in the adult fly brain **(b)** and larval ventral cord **(c)**. GFP1-10 can clearly be seen in cell somas (arrows) and projections (arrowheads). Anti-GFP antibody (cyan) was used to probe for GFP1-10 (cyan), RFP marks Class IV dendritic arborization neuron axon terminals in the GAL4^109-2-80^ > UAS-mCD8::RFP, Actin>GFP1-10 animals, nc82 is a marker of synapses. Scale bar is 50 μm in **b** and 75 μm in **c**.

**Supplementary Figure 3.**
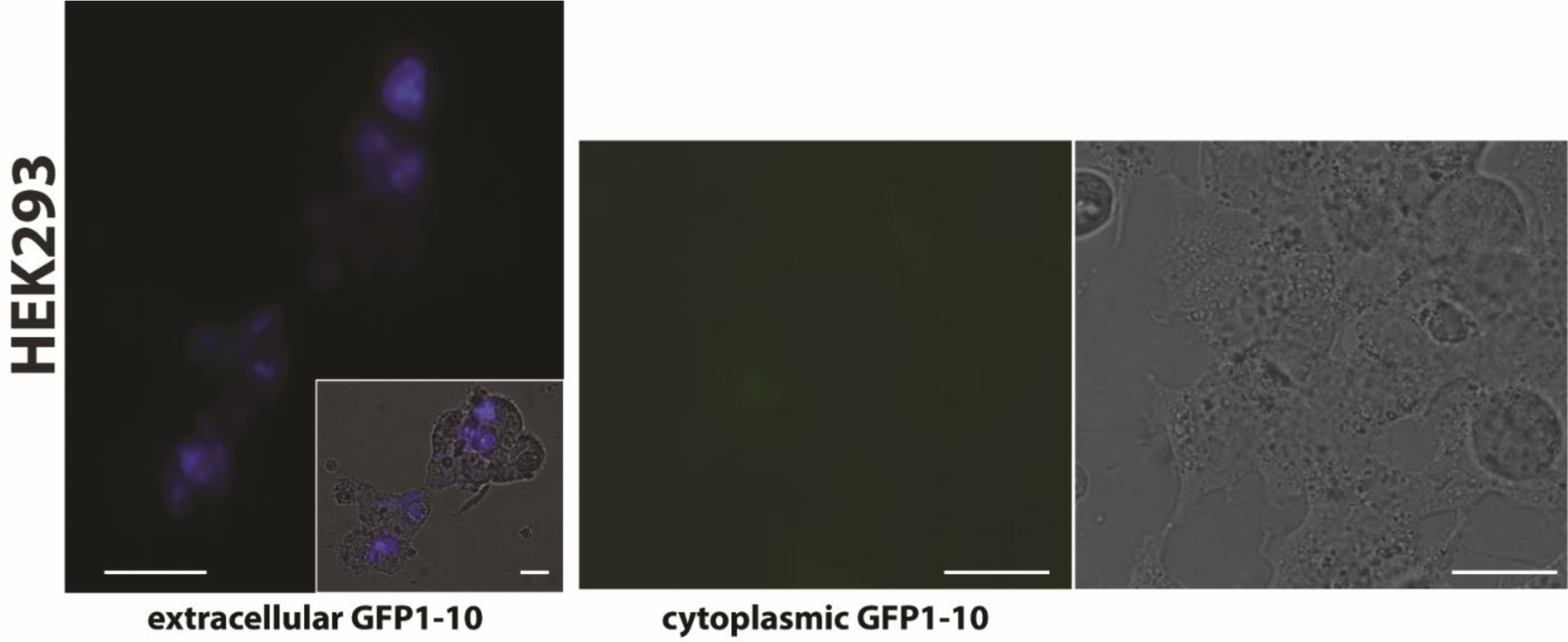
GFP1-10 alone, auto-fluorescence controls. Control images of cells expressing only membrane-bound extracellular expression of GFP1-10 (left images) with Hoechst stain in blue. Cells expressing only cytoplasmic GFP1-10 are shown on the right. Images were acquired at similar excitation and emission settings as experimental data. Scale bars are 10 μm.

**Supplementary Figure 4.**
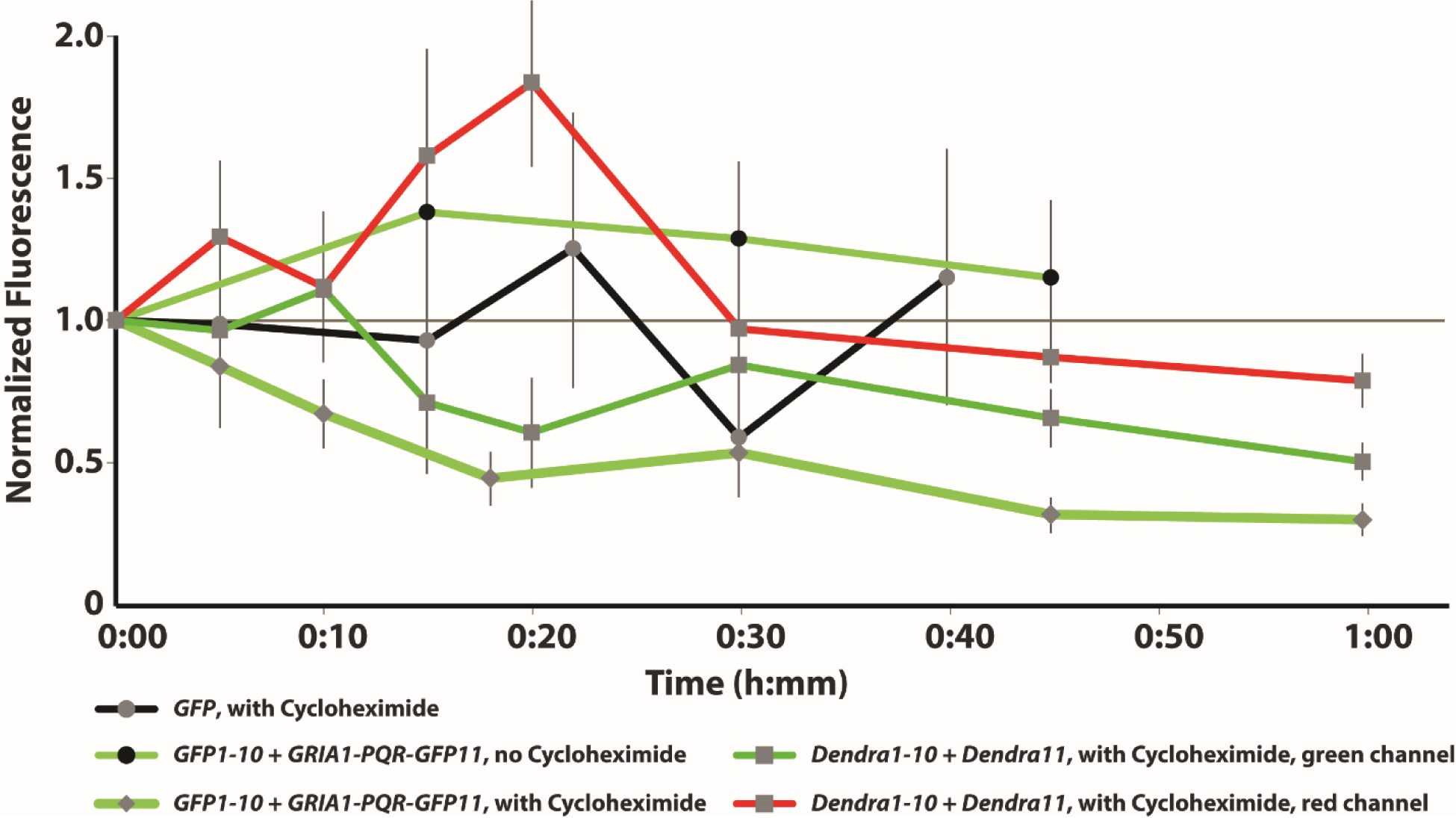
Inhibition of protein synthesis decreases PTR fluorescence. The protein translation inhibitor Cycloheximide was added at time 0:00 to HEK293T cells transfected with *GFP*, PTR with splitGFP, or PTR with splitDendra. Cycloheximide is a small molecule that inhibits translation elongation by binding to the ribosome. Green fluorescence measured in single cells decreased over time after addition of Cycloheximide, but not in cells without Cycloheximide (black circles). The decrease in fluorescence reflected the degradation of the existing fluorescent protein, without a replenishing increase in new protein synthesis when tracked by PTR. Cells transfected with GFP alone (black trace) maintained their overall green fluorescence levels due to the degradation of the existing GFP in addition to the continued production of GFP molecules that were still folding and maturing, which was not the case for cells transfected with PTR constructs. Cells transfected with PTR with splitDendra were photoconverted immediately after time 0:00. All imaging experiments were performed in experimental quadruplicates (*n* > 7 cells per experiment). Error bars are S.D.

**Supplementary Figure 5.**
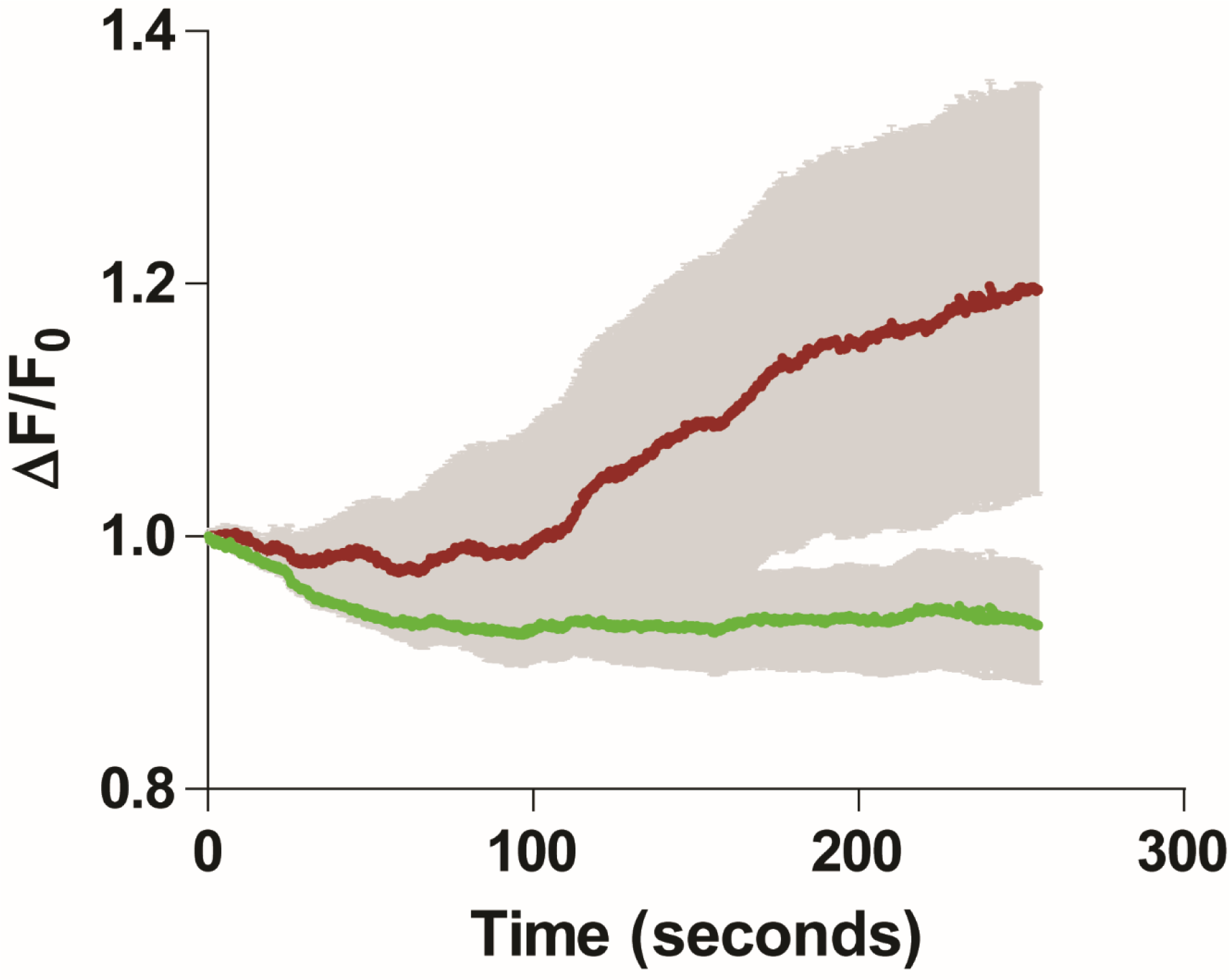
Average GFP fluorescence over time between peri-nuclear and cytoplasmic sites. Reconstituted GFP fluorescence increases over time at peri-nuclear ribosome sites compared to cytoplasmic sites (red versus green traces, respectively; average with standard deviation shown in grey, *n* = 6).

**Supplementary Figure 6.**
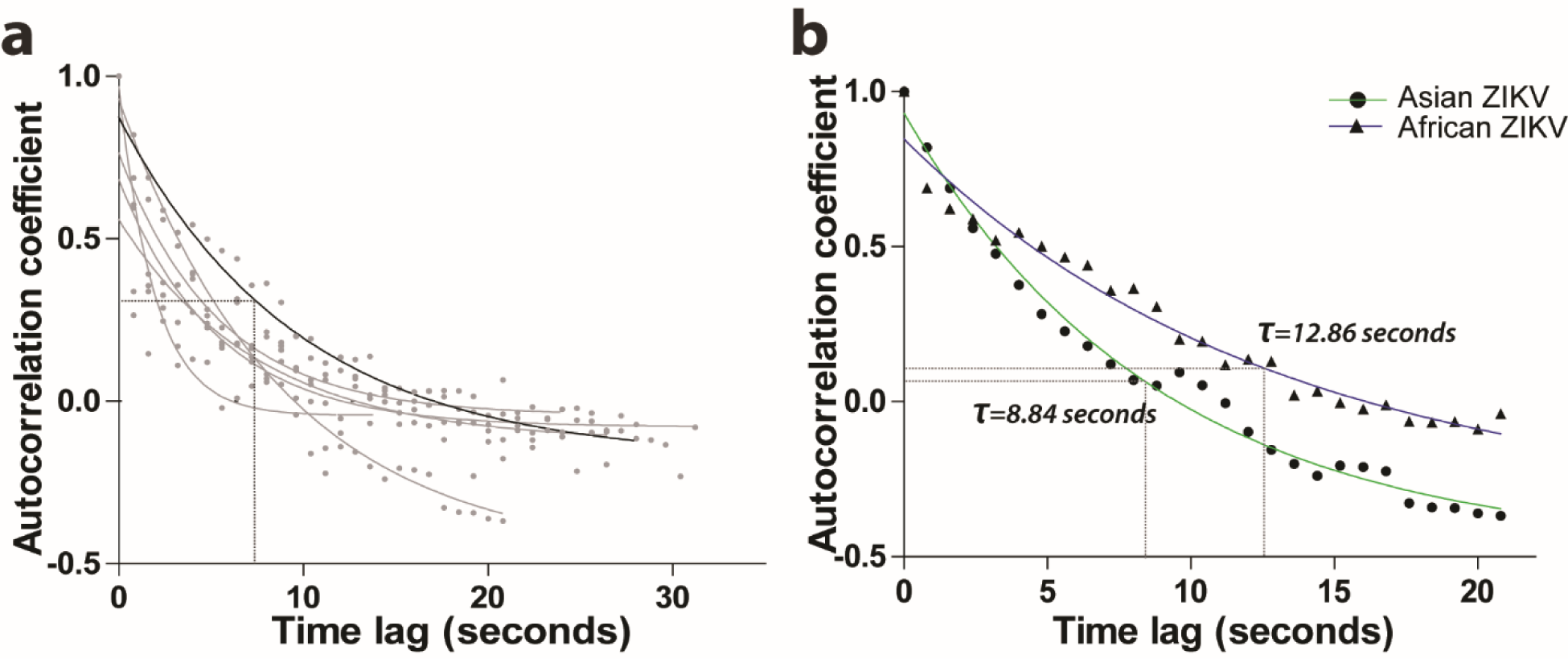
Time course analysis of NS4B protein translation *in vivo*. **a**, Autocorrelation analysis of the fluorescence time courses was used to determine the temporal variation in protein translation burst events between ZIKV strains. **b,** Single exponential decay fits to the autocorrelation function resulted in a time constant of τ = 8.84 ± 9.48 seconds for the Asian ZIKV strain *150989* (*R*^2^ = 0.99, *n* = 3 cells, 2 animals) and τ = 12.86 ± 11.23 seconds for the African ZIKV strain *Ar41524* (*R*^2^ = 0.97, *n* = 4 cells, 2 animals).

**Supplementary Figure 7.**
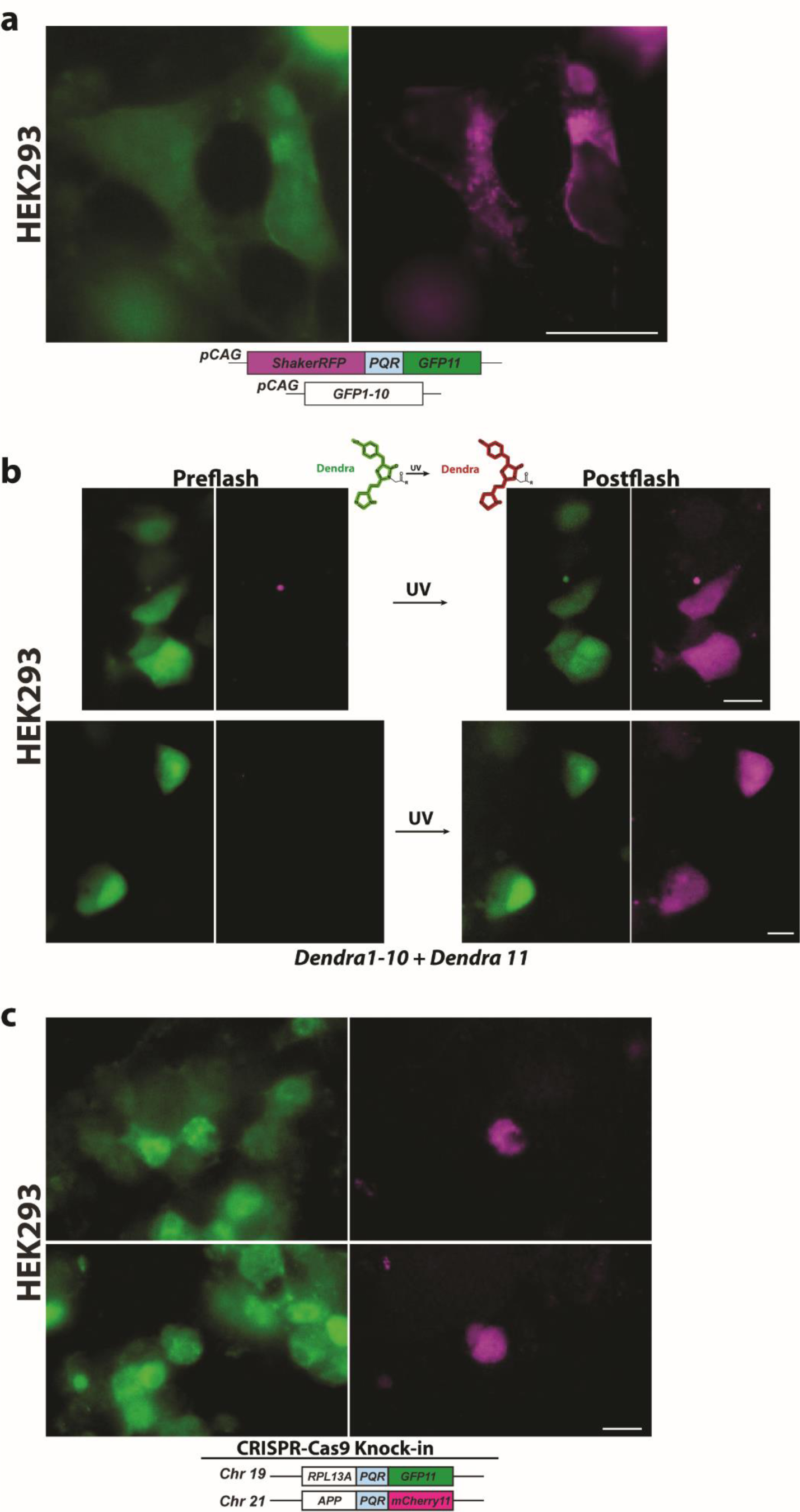
Individual channels for figures. **a**, Green (left) and red (right) channels for **Figure 2a**. Scale bar is 10 μm. **b,** Images of pre-flash UV illumination for both green (left) and red (right) channels for **Figure 3b** are on the far left, and post-flash for green and red channels (left and right images, respectively) are on the far right. Scale bars are 10 μm. **c,** Green (left) and red (right) channels for **Figure 3e**. Scale bar is 10 μm.

**Supplementary Movie Legend 1, Zika Virus Protein Synthesis in Neurons *In Vivo*.**

Time lapse video of neocortical mouse neuron imaged *in vivo* at P23. Fluorescence signal is Zika NS4B protein synthesis, with fluorescence data points from regions of interest taken at 400 millisecond time intervals. Time stamp is seconds with tenths of seconds. Scale is 100 μm on the vertical side.

**Supplementary Movie Legend 2, GluA1 Protein Synthesis in Neurons *In Vitro*.**

Time lapse video of cortical human neuron imaged *in vitro* at 10 days after differentiation. Fluorescence signal is Glutamate Ionotropic Receptor AMPA Type Subunit 1 protein synthesis taken at 25 second time intervals. Time stamp is minutes with seconds. Scale is 100 μm on the vertical side.

**Supplementary Movie Legend 3, Zika Virus Protein Synthesis in Neurons *In Vitro*.**

Time lapse video of cortical human neuron imaged *in vitro* at 10 days after differentiation. Fluorescence signal is Zika NS4B protein synthesis taken at 30 second time intervals. Time stamp is minutes with seconds. Scale is 100 μm on the vertical side.

